# Scaffold protein SHANK3 regulates endothelial cell motility and tissue mechanics

**DOI:** 10.64898/2026.04.12.717721

**Authors:** Megan R. Chastney, Anne Pink, Jouni Härkönen, Gautier Follain, Vanessa Stüve, Joanna W. Pylvänäinen, Anna-Mari Haapanen-Saaristo, Jenna Villman, Monika Vaitkevičiūtė, Giorgio Scita, Illka Paatero, Guillaume Jacquemet, Fabio Giavazzi, Pipsa Saharinen, Johanna Ivaska

**Affiliations:** Turku Bioscience Centre, University of Turku and Åbo Akademi University, Turku, 20520, Finland; Translational Cancer Medicine Program, Research Programs Unit, Biomedicum Helsinki, Haartmaninkatu 8, P.O.B. 63, FI-00014 University of Helsinki, Finland; Department of Pathology, Hospital Nova, Wellbeing Services County of Central Finland, Jyväskylä, Finland; Faculty of Science and Engineering, Cell Biology, Åbo Akademi University, Turku, 20520, Finland; Turku Collegium for Science, Medicine and Technology (TCSMT), Turku, 20520, Finland; InFLAMES Research Flagship, University of Turku, Turku, 20520, Finland; IFOM ETS The AIRC Intitute of Molecular Oncology, Milan, 20139, Italy and University of Milan, Milan, 20100 Italy; Department of Life Technologies, University of Turku, Turku, 20520, Finland; Foundation for the Finnish Cancer Institute, Tukholmankatu 8, Helsinki, 00014, Finland; Department of Medical Biotechnology and Translational Medicine, University of Milan, Milan, 20100 Italy; Wihuri Research Institute, Biomedicum Helsinki, Haartmaninkatu 8, FI-00290 Helsinki, Finland; Department of Biochemistry and Developmental Biology, Faculty of Medicine, University of Helsinki, Finland; Western Finnish Cancer Center (FICAN West), University of Turku, Turku, 20520, Finland

## Abstract

SHANK3 is a multidomain scaffolding protein critical for neuronal function, which has been linked to neurodevelopmental disorders such as autism spectrum disorder. More recently, SHANK3 has been shown to play a role in cell survival and actin dynamics outside the nervous system. Here, we show that SHANK3 is widely expressed in endothelial cells across different tissues, where its role is not well understood. SHANK3 localised to endothelial cell-cell junctions in cultured endothelial cells, and its depletion compromised endothelial barrier function. SHANK silencing altered cell mechanics including elongated cell morphology, reduced cell-matrix traction forces and alteration of cell migration rate. It further triggered dynamic heterogeneity in endothelial monolayers, with regions of coordinated long-range migration interspersed with areas exhibiting only local velocity fluctuations, consistent with a transition toward more fluid-like tissue behaviour. This change in collective dynamics was accompanied by increased spheroid spreading and fusion, suggestive of altered tissue viscosity, and coincided with disrupted cell–cell junction morphology and mechanical forces in SHANK3-depleted cells. *In vivo,* SHANK3 depletion impaired endothelial cell migration, resulting in delayed sprouting of intersegmental vessels and disruption of the vascular network in zebrafish embryos. Furthermore, inducible endothelial-specific deletion of SHANK3 in postnatal mice impaired angiogenic sprouting and reduced vascular complexity in the developing retina. Overall, we demonstrate that SHANK3 plays a role in endothelial cell motility and tissue mechanics, with implications for vascular processes during development.

## Introduction

SH3 and Multiple Ankyrin Repeat Domains 3 (SHANK3) is a large multidomain scaffolding protein that has predominantly been studied in neurons, where it localises to the post synaptic density to regulate neuronal function via varied adaptors, signalling proteins, and the actin cytoskeleton (Duffney et al., 2015; Sarowar & Grabrucker, 2016; Sheng & Kim, 2000). Several mutations in SHANK3 are linked to autism spectrum disorder (ASD), in addition to Phelan-McDermid syndrome and schizophrenia (Durand et al., 2007; Gauthier et al., 2010; Leblond et al., 2014; Moessner et al., 2007; Phelan & McDermid, 2011). SHANK3 is also widely expressed in other cell types and tissues (Lilja et al., 2017), where its varied roles are now being increasingly explored. Beyond neuronal cells, SHANK3 has been shown to regulate cell adhesion via sequestration of the small GTPase Rap1 (Lilja et al., 2017), directly interact with actin to regulate the cytoskeleton (Salomaa et al., 2021), mediate cell survival and proliferative signalling in KRAS mutant cancer cells via interaction with oncogenic KRas (Lilja et al., 2024), and regulate the integrity of the intestinal barrier via tight junction protein ZO-1 (Wei et al., 2017).

Recently, SHANK3 has been shown to be expressed in the endothelial cells of the blood brain barrier where it regulates β-catenin signalling (Y.-E. Kim et al., 2025) and to be upregulated in a subset of lung capillary endothelial cells in response to metastatic melanoma cells (Santio et al., 2024). However, the role of SHANK3 in endothelial cells remains poorly understood.

In addition to maintaining a barrier, endothelial cell migration must also be tightly regulated during tissue homeostasis and vascular development. Angiogenesis, the formation of blood vessels through sprouting from pre-existing vascular segments is a highly regulated process during development, ensuring organ homeostasis. At the cellular level, angiogenesis depends on endothelial cell migration, cell rearrangement, and cell shape changes governed by a complex interplay biochemical and mechanical signals (Angulo-Urarte et al., 2018). Notably, the collective movement of endothelial cells requires the coordination of actin cytoskeleton dynamics with the remodeling of cell-cell junctions such as adherens junctions and tight junctions to maintain vessel integrity during active migration (Paatero et al., 2018).

In this study, we show that SHANK3 is widely expressed in endothelial cells across multiple tissues, and localises to endothelial cell-cell junctions. At this site, SHANK3 is associated with multiple junctional components and actin-regulatory proteins. Depletion of SHANK3 in endothelial cells results in an altered barrier function, reduced cell-ECM forces, and enhanced collective cell migration. Using SHANK3 zebrafish crispants and inducible endothelial SHANK3 deletion in mice, we further demonstrate that SHANK3 is important for angiogenic sprouting during vascular development in the postnatal mouse retina and in zebrafish embryos, highlighting a role for SHANK3 in endothelial cells during vascular mophogenesis.

## Results

### SHANK3 is highly expressed in endothelial cells

Despite predominantly being studied in neurons (and more recently in cancer cells), we found that SHANK3 is also highly expressed in endothelial cells (Fig. 1). Visualisation of single-cell transcriptome data across multiple human tissues and cell types (Tabula Sapiens (Jones et al., 2022)) revealed that SHANK3 is highly expressed in endothelial cells (Fig. 1A), with minimal expression in other cell types. SHANK3 transcripts were detected in endothelial cells from diverse tissues, with the highest levels observed in the spleen, heart, adipose tissue, and trachea (Supplementary Fig. 1A). The lowest expression was observed in endothelial cells in the kidney, large- and small intestine, liver, and the lungs. This variation may reflect endothelial heterogeneity across tissues and potential tissue-specific functions of SHANK3 (Paik et al., 2020; Perez-Gutierrez et al., 2024). Additionally, SHANK3 mRNA expression was observed across multiple endothelial cell types from different vascular beds, including capillary, venous, arterial and lymphatic endothelial cells (Supplementary Fig. 1B). Gene set enrichment analyses (GSEA) showed several positively enriched gene ontology (GO) terms in cells with high SHANK3 mRNA expression that were related to vascular development (e.g. endothelium development, circulatory system development, vasculature development, and blood vessel morphogenesis) (Supplementary Fig. 1C). Additionally, multiple GO terms related to previously recognised roles for SHANK3 were also identified, including actin-related terms (actin binding, regulation of actin filament organisation, and actin filament binding) and cell junctions (cell junction organisation) (Y.-E. Kim et al., 2025; Salomaa et al., 2021; Sarowar & Grabrucker, 2016; Wei et al., 2017). We also detected SHANK3 expression in CD31 (PECAM-1)-positive endothelial cells in diverse human tissues, including capillaries, veins and arteries in the lung (Fig. 1B), as well as in the vasculature in the appendix, kidney, and soft tissue (Supplementary Fig. 1D).

**Figure 1.**
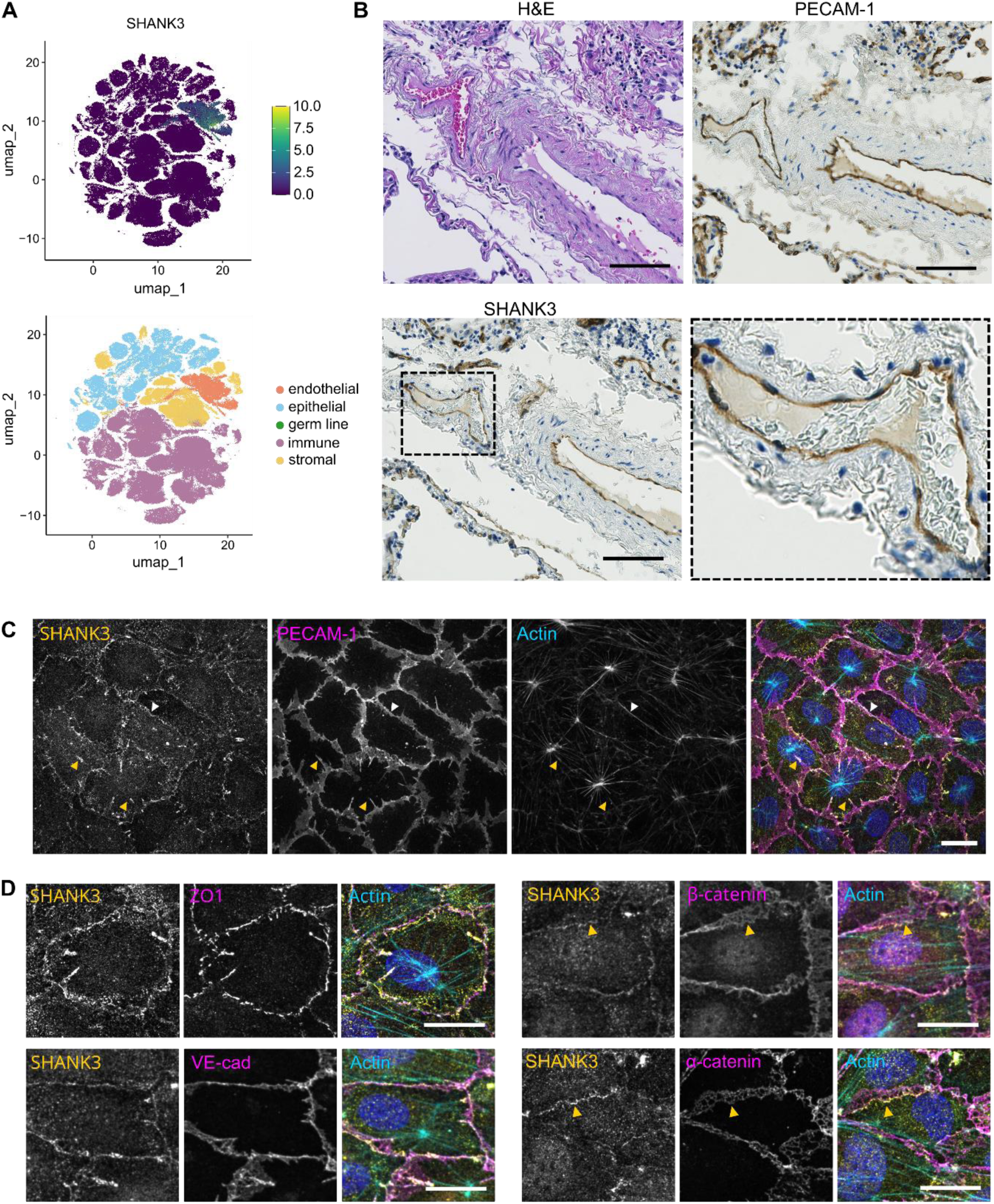
SHANK3 is highly expressed in endothelial cells and localises to cell junctions. **A.** UMAP of SHANK3 mRNA expression from single-cell transcriptome data (Tabula Sapiens (Jones et al., 2022)) shows high expression in endothelial cells. **B.** Histological staining of serial sections of human lung tissue. Insert shows SHANK3 expression in vein endothelial cells. PECAM-1 used as endothelial cell marker. Scale bar 100 µm. H&E, hematoxylin and eosin. **C.** Immunofluorescence images of HUVEC monolayers showing SHANK3 at linear cell junctions (white arrows) and protrusive structures that link to actin (orange arrows). Scale bar 20 µm. **D.** SHANK3 colocalises with cell junction components ZO-1, VE-cadherin (VE-cad) β-catenin and α-catenin. Yellow arrows indicate reticular adherens junctions. Scale bar 20 µm.

### SHANK3 localises to cell junctions in endothelial cells

To determine the subcellular localisation of SHANK3 in endothelial cells, we stained for endogenous SHANK3 in Human Umbilical Vein Endothelial Cell (HUVEC) monolayers (Fig. 1C). SHANK3 localised to cell junctions, with PECAM-1 used as a junctional marker. SHANK3 was present both in linear junctions with parallel cortical actin, resembling linear adherens junctions typically associated with mature, stable cell junctions (Huveneers et al., 2012) (Fig. 1C, white arrows) and appeared particularly enriched at finger-like protrusive structures that linked actin-stress fibres in neighbouring cells connecting to central ‘aster’-like actin structures (orange arrows). These actin aster-like structures lay at the basal side of the cell, with radial actin fibres connecting either SHANK3-positive cell junctions, or paxillin positive integrin adhesion complexes (IACs) at the ventral surface (Supplementary Fig. 2A). Similar structures have previously been observed in cultured arterial endothelial cells (van Geemen et al., 2014), and could also resemble the highly organised supracellular actin network in intestinal villi epithelial cells that provides mechanical stability to the tissue (Barai et al., 2025). Due to the centralised location and elongated morphology of the actin-associated IACs, we tested whether these adhesions represented fibrillar adhesions, a specialised subtype of IAC that facilitate fibronectin fibrillogenesis (Pankov et al., 2000; Zamir et al., 2000). Notably, these structures were enriched in fibrillar adhesion markers tensin-1 and integrin α5, and associated with fibronectin fibres (Supplementary Fig. 2B), supporting their involvement in fibronectin fibrillogenesis in endothelial cells. Additionally, phospho-Myosin Light Chain (pMLC) staining indicated that these aster-like structures are highly contractile, displaying a prominent contractile core (Supplementary Fig. 2C, green arrows) and elevated, focalised pMLC at the termini of radial arms, at protrusive cell-junctions (Supplementary Fig. 2C, yellow arrows). Live cell imaging of actin in HUVECs expressing GFP-CAAX as a membrane marker revealed that these structures were relatively stable in relation to the ECM, likely because of their anchorage to the ECM via the fibrillar adhesions (Supplementary Video 1). The protrusive structures resembled engulfed cadherin fingers, polarised junctional structures that mediate directional migration in endothelial cell monolayers (Hayer et al., 2016). However, unlike canonical cadherin fingers, the structures observed here often lacked the coordinated cell-wide alignment required to support directional migration.

Endothelial cells contain multiple cell junction subtypes, including tight junctions (containing claudins, occludin, and ZO-1) and adherens junctions (including JAMs, PECAM-1, Vascular Endothelial (VE)- cadherin and α- and β-catenin) (Dejana, 2004). Co-staining of SHANK3 with several junction markers revealed co-localisation with VE-cadherin, α-catenin and β-catenin (Fig. 1D). However, co-localisation was particularly prominent with ZO-1, especially at protrusive junctions. Although the adaptor protein ZO-1 is found at both tight junctions and adherens junctions, the presence of SHANK3 at only one edge of reticular adherens junctions observed with α-catenin and β-catenin staining (Fig. 1D, yellow arrows) suggests that SHANK3 may preferentially associate with more apically-localised junctional domains (Fernández-Martín et al., 2012).

### The SHANK3 proximity interactome identifies actin- and cell junction-associated components

To identify potential SHANK3 regulatory proteins, we performed a proximity biotinylation interactome screen using BioID (D. I. Kim et al., 2014). To maximise the number of proximity interactors and capture potential N- or C- terminal domain specific interactors, SHANK3 was fused with the promiscuous biotin ligase, BirA* (BirA), at either the N- or C-terminus (BirA-SHANK3 and SHANK3-BirA, respectively) and stably expressed in U2OS cells, with BirA alone serving as a negative control. While BirA control showed no specific subcellular localisation, BirA-SHANK3 and SHANK3-BirA localised to the cell membrane, including lamellipodia and cell junctions (Fig. 2A). Proteomic analysis of biotinylated proximity interactors revealed 87 high confidence (BFDR ≤ 0.05) BirA-SHANK3 interactors and 47 SHANK3-BirA interactors (94 in total, Fig. 2B and Supplementary Table 1). There was a large overlap in proximity interactors identified by the N- and C-terminally tagged BirA-SHANK3 and SHANK3-BirA (41). Mapping of biotinylated peptides in SHANK3 identified by both constructs revealed an overlap in biotinylation in the central domains of SHANK3, suggesting that with our strategy of tagging SHANK3 at both termini the BirA tag is able to label a significant proportion of SHANK3, and therefore also likely to label SHANK3 interactors spanning the entire length of this large scaffolding protein (Supplementary Fig. 3A).

**Figure 2.**
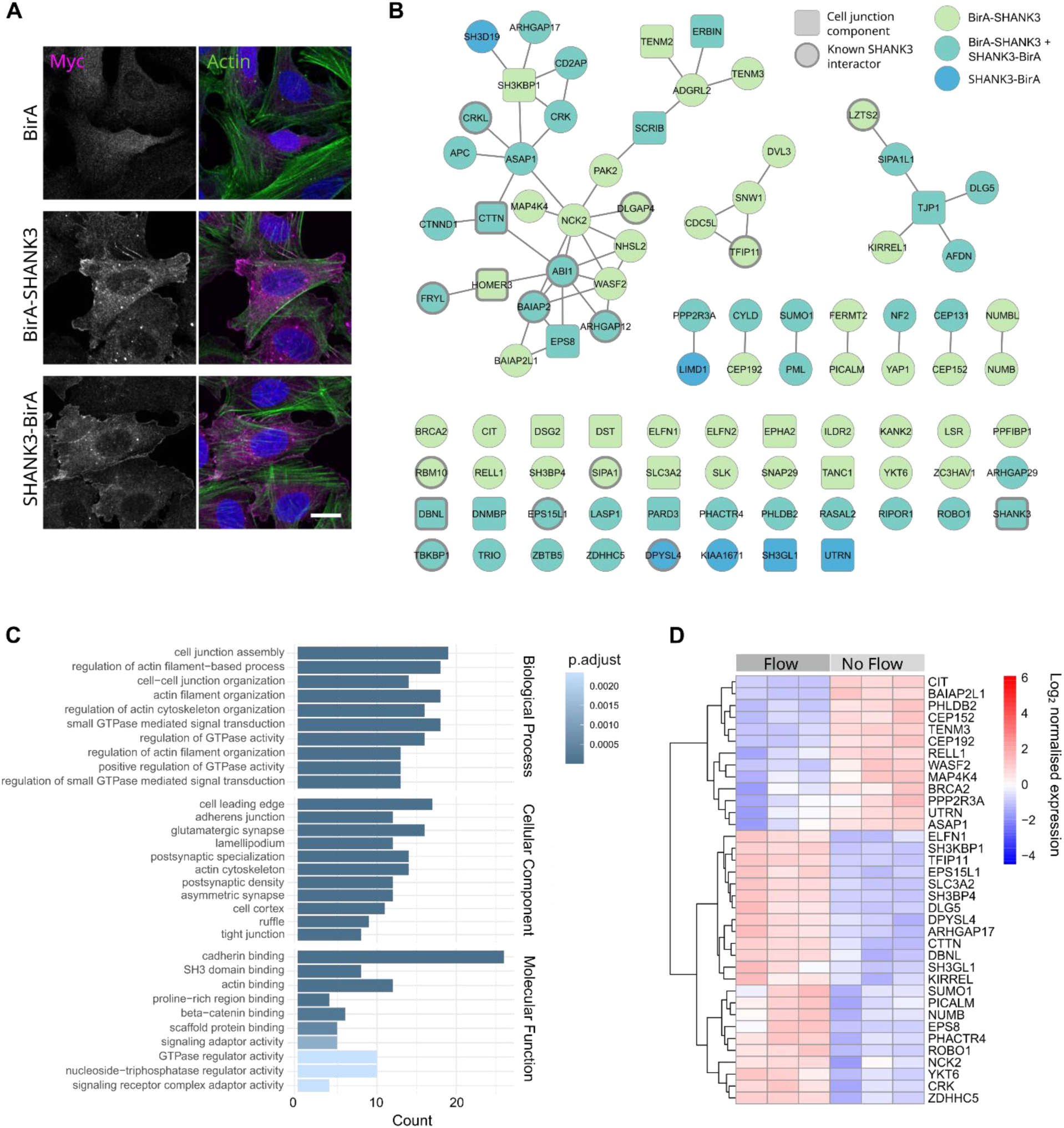
SHANK3 proximity biotinylation reveals actin- and cell junction-associated proximity interactors. **A.** Immunofluorescence images showing subcellular localisation of myc-tagged BirA, BirA-SHANK3 and SHANK3-BirA in stably-expressing U2OS cells. Scale bar 20 µm. **B.** Interaction network of SHANK3 proximity interactors (BFDR ≤ 0.05). Nodes correspond to proximity interactors and edges indicate protein-protein interactions from STRING-db (Szklarczyk et al., 2023). BirA-SHANK3 interactors shown in green, SHANK3-BirA shown in blue, and shared interactors shown in turquoise. Square nodes indicate cell junction components (UniProt keyword ‘cell junction’; KW-0965) and thick outlines indicate previously identified SHANK3 interactors (BioGRID). **C.** GO analysis of SHANK3 proximity interactors (BFDR ≤ 0.05). The top 10 terms in each category are shown. **D.** SHANK3 proximity interactors with mRNA expression significantly up or down regulated in HUVECs in the presence or absence of laminar flow (Follain et al., 2021).

Consistent with the observed localisation of SHANK3 at cell junctions and its previous known role in regulating the actin cytoskeleton, 29 of the proximity interactors (30.9 %) were annotated as cell junction proteins (UniProt key word ‘Cell junction’; KW-0965), and 24 (25.5 %) as cytoskeletal proteins (UniProt key word ‘Cytoskeleton’; KW-0206). For example, we identified several cell-cell junction and actin adaptors involved in endothelial junctions, including ZO-1 (TJP1), p120-catenin (CTNND1), afadin (AFDN), and EPS8 (Birukova et al., 2012; Dejana, 2004; Giampietro et al., 2015). Additionally, 16 proteins (17 %) identified in the screen have previously been reported as SHANK3 interactors (BioGRID), including cortactin (CTTN), IRSp53 (BAIAP2) and ableson interactor 1 (ABI1). Gene Ontology (GO) analysis of the SHANK3 proximity interactome revealed enrichment of terms related to cell junctions and actin regulation (Fig. 2C). GO terms linked to GTPase signalling were also identified, consistent with the reported role of SHANK3 in regulating Rap/Ras signalling in cancer cells (Lilja et al., 2017, 2024). Selected BioID hits, together with previously reported SHANK3 interactors not detected in the proximity interactome (SHARPIN, β-catenin), were further validated in U2OS cells using a ‘knock sideways’ approach, in which SHANK3-mCherry-zdk was redirected to mitochondria via TOMM20-LOV2 (Supplementary Fig. 3B and C). Antibodies were used to detect endogenous targets, and overlapping co-localisation with SHANK3-mCherry-zdk (but not mCherry-zdk alone) suggested an interaction between the two proteins in the cell. The junctional protein ZO-1, actin regulatory protein WAVE2 (WASF2), the ARF-GAP ASAP1, and the Bar-domain containing protein IRSp53 were recruited to mitochondria together with SHANK3 (Supplementary Fig. 3B). Conversely, other proteins, such as EPHA2 and α-catenin, did not relocalise to mitochondria with SHANK3-mcherry-zdk (Supplementary Fig. 3C), indicating the specificity of our assay. Although not detected in the BioID dataset, the known SHANK3 interactors SHARPIN and β-catenin, were included as positive controls and showed robust colocalization with SHANK3-mCherry-zdk.

Additionally, 36 out of 94 of the proximity interactors (38.3%) were previously found to be significantly up or down regulated in endothelial cells (HUVECs) in response to laminar flow (Fig. 2D) (Follain et al., 2021), suggesting that these proteins are involved in endothelial cell biology. Overall, the SHANK3 proximity interactome provides a useful resource for further exploration of SHANK3 functions.

### Depletion of SHANK3 alters endothelial cell morphology, barrier function, and migration

To examine the effects of SHANK3 depletion on endothelial cells, we knocked down SHANK3 using two previously validated siRNAs targeting SHANK3 (Lilja et al., 2024) (Fig. 3A). While control HUVECs (AllStars siRNA non-targeting control), typically exhibited a regular polygonal morphology within the monolayer, SHANK3 knockdown resulted in an increased proportion of elongated cells, characterised by reduced circularity and increased aspect ratio (Fig. 3B and C, Supplementary Fig. 4A). This observation was validated in HUVECs stably expressing doxycycline-inducible shRNA directed against SHANK3 (shSHANK3 cells) (Supplementary Fig. 4B-D). Furthermore, depletion of SHANK3 also led to increased gaps in the HUVEC monolayer (Fig. 3D and E, Supplementary Fig. 4E), as observed by increased accessibility of the fibronectin antibody to the underlying extracellular matrix in non-permeabilised fixed cells (Ball et al., 2024). This indicates that SHANK3 plays a role in maintaining barrier integrity in endothelial cells.

**Figure 3.**
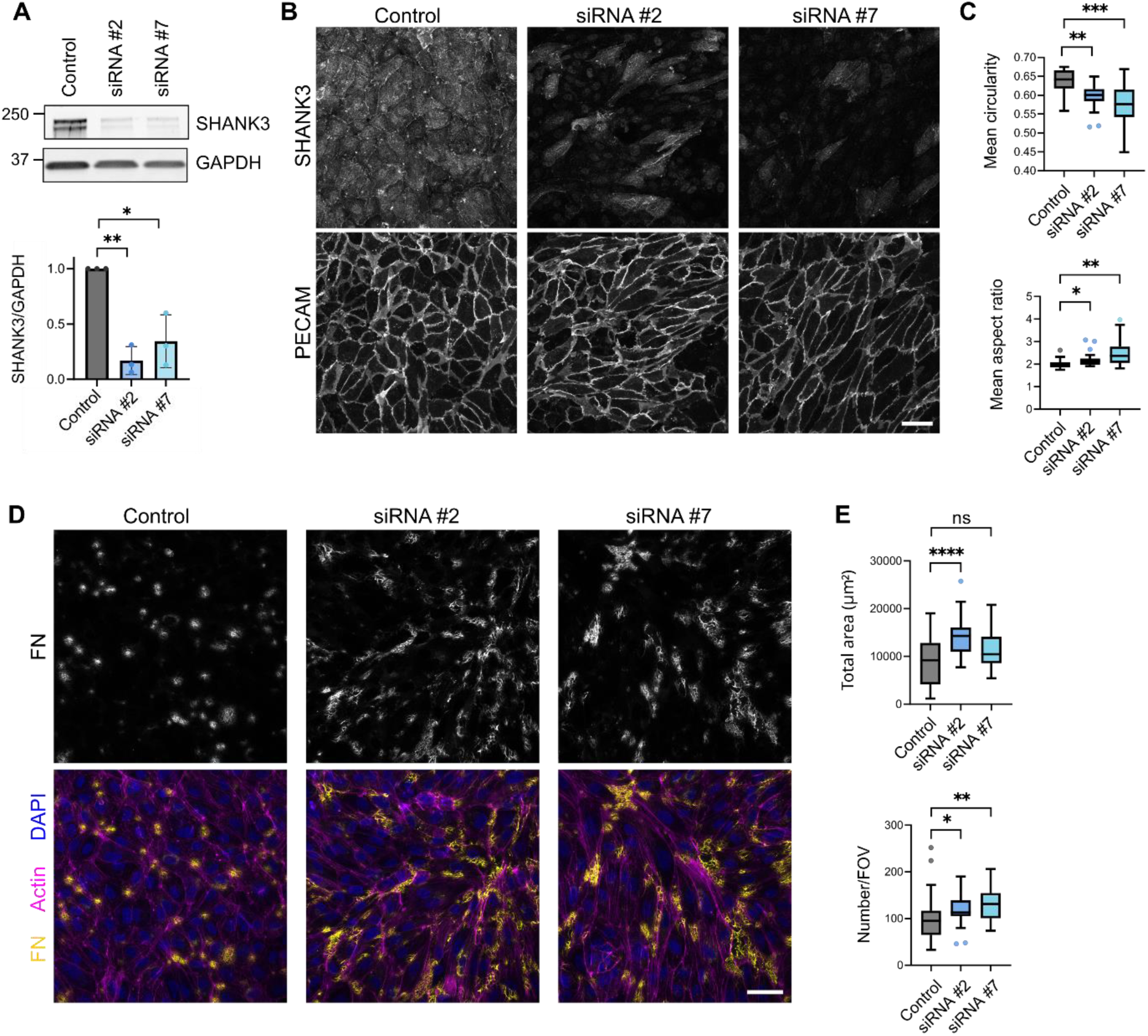
SHANK3 depletion alters cell morphology and affects barrier function. **A.** Representative western blot and quantification of SHANK3 depletion in HUVECs using two siRNAs targeting SHANK3. GAPDH used as a loading control. Paired t-test. n = 3 biological replicates. **B.** Immunofluorescence images of HUVEC monolayers following siRNA depletion of SHANK3, showing SHANK3 depletion and altered cell morphology. Scale bar 50 µm. **C.** Quantification of cell shape from B using PECAM-1 to identify cell boundaries. Means per field of view shown. n = 3 biological replicates, 6 fields of view per condition each replicate. Unpaired t-test. **D.** Detection of gaps in the HUVEC monolayer using anti-fibronectin antibodies in non-permeabilised cells. Scale bar 50 µm. **E.** Quantification of fibronectin patches in D. n = 3 biological replicates, 10-13 fields of view per condition each replicate. Unpaired t-test. Control, AllStars negative control; ns, non-significant; ****, p value < 0.001; ***, p value < 0.005; **, p value < 0.01; *, p value < 0.05.

Changes in cell shape within cell monolayers are indicative of altered collective migration and tissue mechanical states (Atia et al., 2018; Bi et al., 2015; Blauth et al., 2021; Park et al., 2015). In particular, increased cell elongation has been associated with the transition from a jammed, solid-like state to a more fluid-like, unjammed state characterised by an increased rate of cell rearrangement events. To determine whether the morphological changes observed upon SHANK3 depletion were accompanied by altered cell motility, we tracked the motion of cell nuclei over 16 h in SHANK3-depleted HUVEC monolayers (Fig. 4A). SHANK3-depleted shRNA cells showed an increased migration speed, total distance travelled and track displacement increased over controls (Fig. 4A and B). No significant difference in wound closure rates were observed in a scratch wound assay (Supplementary Fig. 5A and B), suggesting that the changes in migration may be specific to cell monolayers rather than collective- or individual cell migration into cell-free spaces. The study of the two-point velocity correlation function revealed only a weak mutual alignment between adjacent cells in all conditions, in the absence of any long-range coordinated collective migration (Fig. 4C, Supplementary Fig. 5C). Similarly, SHANK3 depletion did not increase the frequency of local cell rearrangements (T1 events per unit time), suggesting that the overall fluidity of the monolayer was not substantially altered (Fig. 4D, E, Supplementary Fig. 5D).

**Figure 4.**
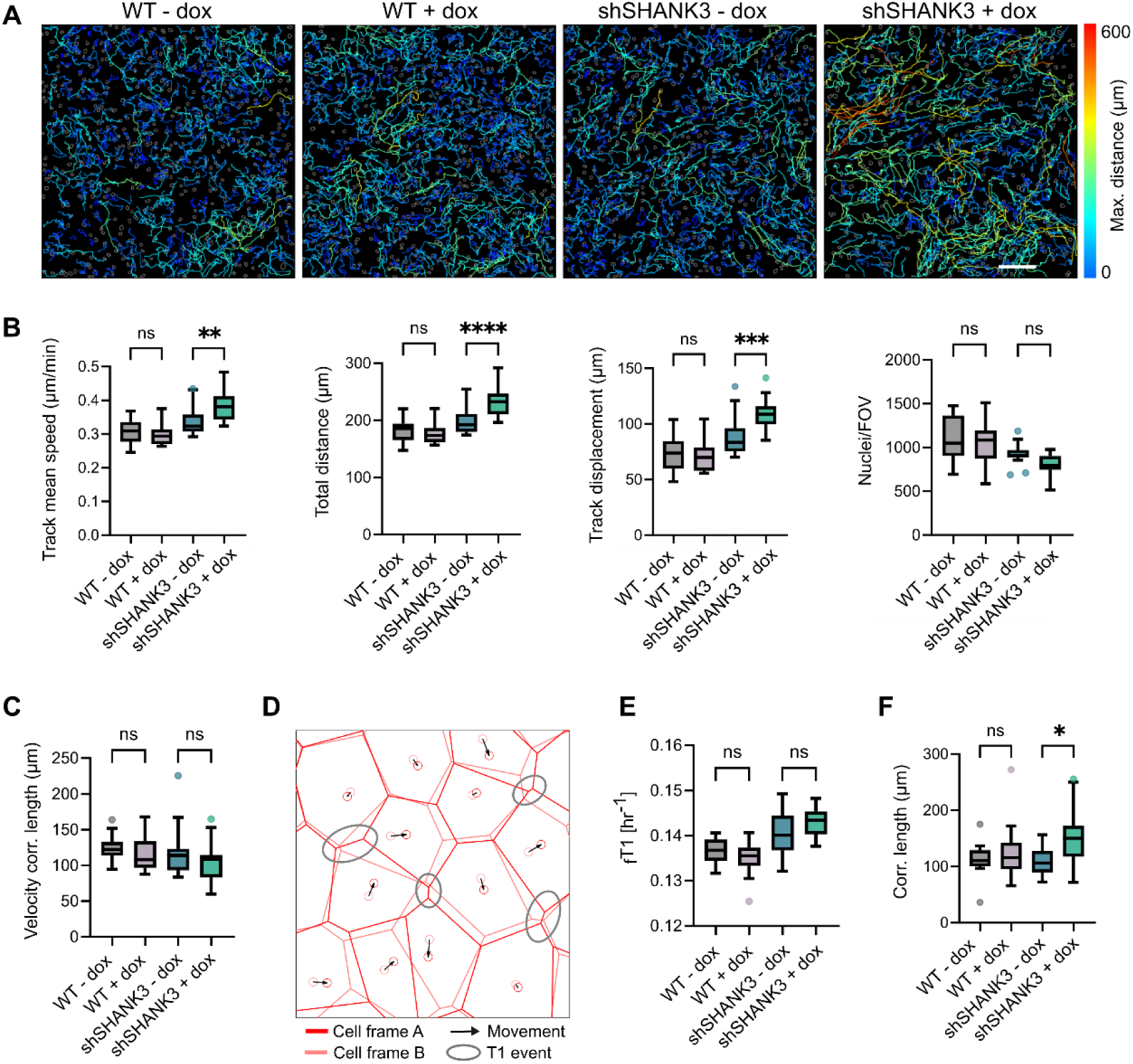
SHANK3 depletion leads to increased cell migration in endothelial cell monolayers. **A.** Migration of WT and shSHANK3 HUVECs +/− doxycycline (dox) in a monolayer. Nuclei were segmented (grey outlines) and tracked every 10 minutes over 16h. Max distance of tracks shown. Scale bar = 200 µm. **B.** Quantification of migration parameters and nuclei count from A. n = 3 biological replicates, 4-6 fields of view per condition each replicate. One-way ANOVA with Šídák test for multiple comparisons used for statistical analysis. **C**. Velocity correlation length obtained from the two-point velocity correlation function 〈𝒏(𝒙) ⋅ 𝒏(𝒙 + 𝒓)〉 for WT and shSHANK3 HUVECs +/− dox. n = 3 biological replicates, 5 fields of view per condition in each replicate. One-way ANOVA with Šídák test for multiple comparisons used for statistical analysis. **D.** Diagram showing cell rearrangements in monolayers (T1 events) between two timepoints. Pink and red indicate cell boundaries at timepoint A and B, respectively. Grey circles indicate switching of cell neighbours between timepoints. Arrows indicate direction of movement. **E.** Mean frequency of T1 events per cell (f_T1_) from WT and shSHANK3 HUVECs +/− dox in monolayers. n = 3 biological replicates, 5 fields of view per condition in each replicate. One-way ANOVA with Šídák test for multiple comparisons used for statistical analysis. **F.** Correlation length obtained from the two-point correlation function in the root mean square velocity 〈𝑣^2^(𝒙)𝑣^2^(𝒙 + 𝒓)〉.n = 3 biological replicates, 5 fields of view per condition in each replicate. One-way ANOVA with Šídák test for multiple comparisons used for statistical analysis. ns, non-significant; ****, p value < 0.001; ***, p value < 0.005; **, p value < 0.01; *, p value < 0.05.

However, analysis of two-point correlations in the root mean square velocity revealed increased spatial segregation between highly motile and less motile cells in SHANK3-depleted monolayers, indicating the emergence of spatially heterogeneous dynamics (Fig. 4F, Supplementary Fig. 5E). Such dynamic heterogeneity is a distinctive feature of disordered systems approaching kinetic arrest or undergoing a rigidity transition (Berthier & Biroli, 2011; Park et al., 2015).

The transition between jammed and unjammed states in epithelial tissues has been shown to be regulated by a variety of biomechanical and signalling cues. For example, hyperactivation of ERK signalling, resulting from overexpression of the endocytic trafficking regulator RAB5A, can reawaken collective cell migration in otherwise jammed epithelial monolayers (Palamidessi et al., 2019). SHANK3 has previously been identified as a direct inhibitor of Ras–Raf signalling, and its depletion leads to hyperactivation of ERK signalling in KRas-mutant cancer cells (Lilja et al., 2024). We therefore tested whether SHANK3 depletion affects ERK signalling and ERK-mediated migration in HUVECs (Supplementary Fig. 5F-H). In contrast to KRas-mutant cancer cell lines, no differences in ERK activity were observed between SHANK3-silenced and control HUVECs, likely reflecting the absence of oncogenic KRas signalling in these cells (Supplementary Fig. 5F, G). However, pharmacological inhibition of ERK signalling using the MEK inhibitor trametinib reduced HUVEC migration velocity in both control and SHANK3-depleted cells, and restored the elevated migration speed of SHANK3 knockdown cells to levels comparable with controls (Supplementary Fig. 5H). Together, these results indicate that while ERK signalling contributes to endothelial cell motility, hyperactivation of ERK signalling is unlikely to be the primary driver of the increased migration observed following SHANK3 depletion.

### SHANK3 deficient endothelial cells show altered viscosity and dynamics

To test whether the differences in migration between control and SHANK3-depleted HUVECs affect their migration under more physiological conditions, we performed a tubule formation assay, in which HUVECs were plated onto a basement membrane-like substrate (Matrigel) and allowed to migrate and form capillary-like tubules, mimicking early angiogenic morphogenesis (Fig. 5A and Supplementary Fig. 6A-C). Despite the changes in migration observed in monolayers, no significant differences were detected in tubule formation metrics, including mean mesh size, mesh index, and branching interval (Supplementary Fig. 6A, B). However, clear differences in tubule morphology were observed between SHANK3-depleted and control HUVECs. Immunofluorescence imaging of the tubules revealed that SHANK3-depleted HUVECs have uneven, ‘lumpy’ edges compared to the smooth edges of the control HUVEC tubules (Fig. 5A). This was quantified by comparing the baseline of the edge with the traced edge to give an ‘edge score’, with SHANK3-depleted cells having a significantly higher edge score (Fig. 5B, C).

**Figure 5.**
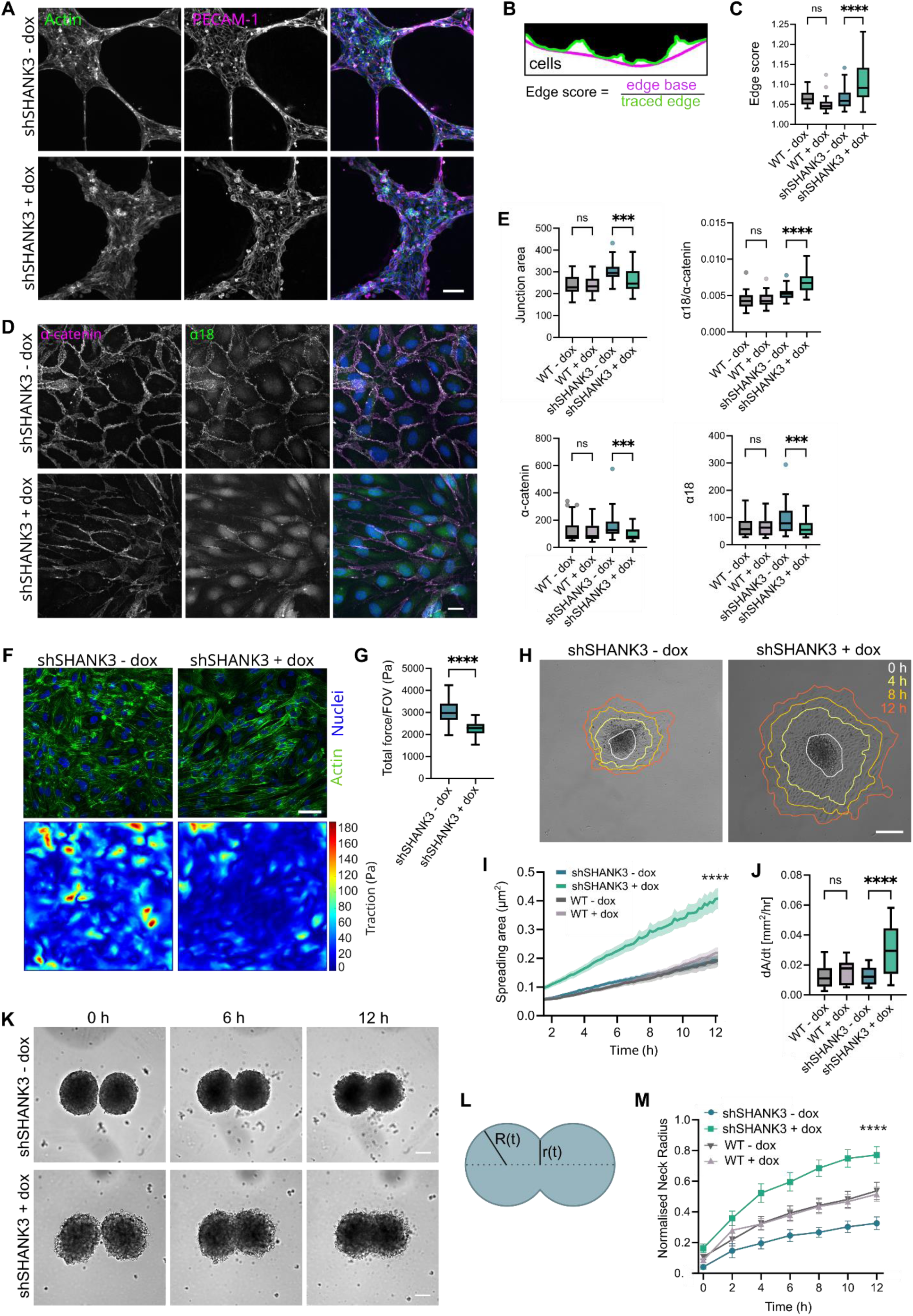
SHANK3-depleted endothelial cells show altered tissue dynamics. **A.** Immunofluorescence imaging of shSHANK3 HUVECs +/− doxycycline (dox) seeded onto Matrigel for 16 h. Scale bar 200 µm. **B.** Schematic of ‘edge score’ used to score HUVEC morphology in A. **C.** Quantification of edge score from A and supplementary Fig. 6C. One way ANOVA with Šídák test for multiple comparisons used for statistical analysis. n = shSHANK3 cells, 3 independent experiments; WT cells, 1 experiment; 15-24 edges quantified per replicate. **D.** Immunofluorescence images of shSHANK3 HUVEC monolayers +/− dox stained for total α-catenin and force-sensitive α-catenin conformation (α18). Scale bar 20 µm. **E.** Quantification of cells from D and Supplementary Fig. 6D. One-way ANOVA with Šídák test for multiple comparisons used for statistical analysis. n = 4 biological replicates, 9-11 fields of view quantified per replicate. α18/α-cat IntDen is normalised to nuclei per field of view. **F.** Traction force microscopy of shSHANK3 HUVECs +/− dox monolayers. Scale bar 50 µm. **G.** Quantification of total traction force per field of view (Pa) from F. n = 27-29 fields of view per condition across two independent experiments. **H.** Spheroid wetting assay tracking shSHANK3 HUVEC +/− dox migration away from spheroid over 12 h. Overlay indicates cell periphery at indicated time points. Scale bar 200 µm. **I.** Quantification of cell outgrowth area over time in H and Supplementary Fig. 6G. One way ANOVA with Šídák test for multiple comparisons used for statistical analysis at end timepoint. n = 16-24 spheroids per condition over three independent experiments. **J.** Spheroid spreading rate (dA/dt [mm^2^/hr]) of spheroids from H and Supplementary Fig. 6G. n = 16-24 spheroids per condition over three independent experiments. One way ANOVA with Šídák test for multiple comparisons used for statistical analysis. **K.** Spheroid merge assay showing merging of shSHANK3 HUVEC spheroids +/− dox over 12 h. Scale bar 100 µm. **L.** Diagram of measurements of spheroids taken to calculate normalised neck radius in M. **M.** Quantification of normalised neck radius (R(t)/r(t)) of spheroids in K and Supplementary Fig. 6H. n = 4 biological replicates WT +/− dox and shSHANK3 + dox and 3 biological replicates shSHANK3 - dox, 2-6 spheroids per replicate. One-way ANOVA with Šídák test for multiple comparisons used for statistical analysis at 12 h. FOV, field of view; ns, non-significant; ****, p value < 0.001; ***, p value < 0.005.

We hypothesised that this altered edge morphology may result from changes in forces at cell junctions or traction against the substrate. As a read out of forces at cell junctions, we used an antibody directed against a force-sensitive epitope within α-catenin (α18) (Fig. 5D) (Yonemura et al., 2010). This showed that overall junction area (measured by masked α-catenin) was decreased in SHANK3 suggesting that junctions may be impaired (Fig. 5E). However, within these junctions, the ratio of α18 compared to α-catenin was higher, suggesting that they were under higher strain, possibly compensating for the lower cell junction area per cell. Furthermore, using traction force microscopy we showed that SHANK3 depletion reduced traction forces on the substrate (Fig. 5F,G and Supplementary Fig. 6F). Thus, in the absence of SHANK3 the disruption in cell junctions and traction forces may affect the ability of cells to resist tensile forces that ordinarily lead to the smooth edges seen in control cells.

Mechanical forces at cell–cell and cell-ECM junctions contribute to the effective viscosity of epithelial tissues. One experimental approach to probe tissue viscosity is the spheroid wetting assay, in which multicellular spheroids are placed onto a substrate and allowed to spread, or “wet”, the surface (Douezan et al., 2011; Pérez-González et al., 2019). The kinetics of spheroid spreading depend on several factors, including cellular contractility, cell–ECM adhesion, and the strength of cell–cell junctions. In SHANK3-depleted cells, spreading velocity was significantly increased compared with control cells (Fig. 5H-J, Supplementary Fig. 6G), suggesting a lower viscosity.

Consistent with the altered morphology observed in cells plated onto Matrigel, SHANK3-depleted spheroids also exhibited less well-defined edges compared with control spheroids. To further assess tissue mechanical properties, we performed a spheroid-fusion assay, in which two spheroids merge over time, analogous to the coalescence of viscoelastic droplets (Dechristé et al., 2018; Marchesi et al., 2026). In this assay, spheroid fusion rate reflects the ratio between tissue spheroid surface tension and viscosity. SHANK3-depleted spheroids fused more rapidly than control spheroids, as indicated by an increased normalised neck radius during the fusion process (Fig. 5K-M, Supplementary Fig. 6H). Together, these results indicate that depletion of SHANK3 induced an alteration in the mechanical properties of HUVEC aggregates, consistent with a reduction of the effective tissue viscosity.

### SHANK3 is required for efficient angiogenesis during development

To test whether SHANK3 plays a role in cell migration in more physiological conditions, we examined the effects of SHANK3 depletion on endothelial cells in 3D cell culture. Sprouting of cells from spheroids in 3D collagen I gels revealed that although the average length of sprouts per spheroid was not significantly different, SHANK3-depleted cells had a higher number of smaller sprouts (< 100 µm) than control cells, and had very few longer sprouts (> 100 µm) (Fig. 6A, B). This could indicate that while initial sprout formation is unaffected, coordination of the cell migration is impaired, perhaps through the altered cell-junctions observed in 2D monolayers. We therefore explored whether SHANK3 is involved in vascular development in zebrafish embryos and in the postanatal mouse retina.

**Figure 6.**
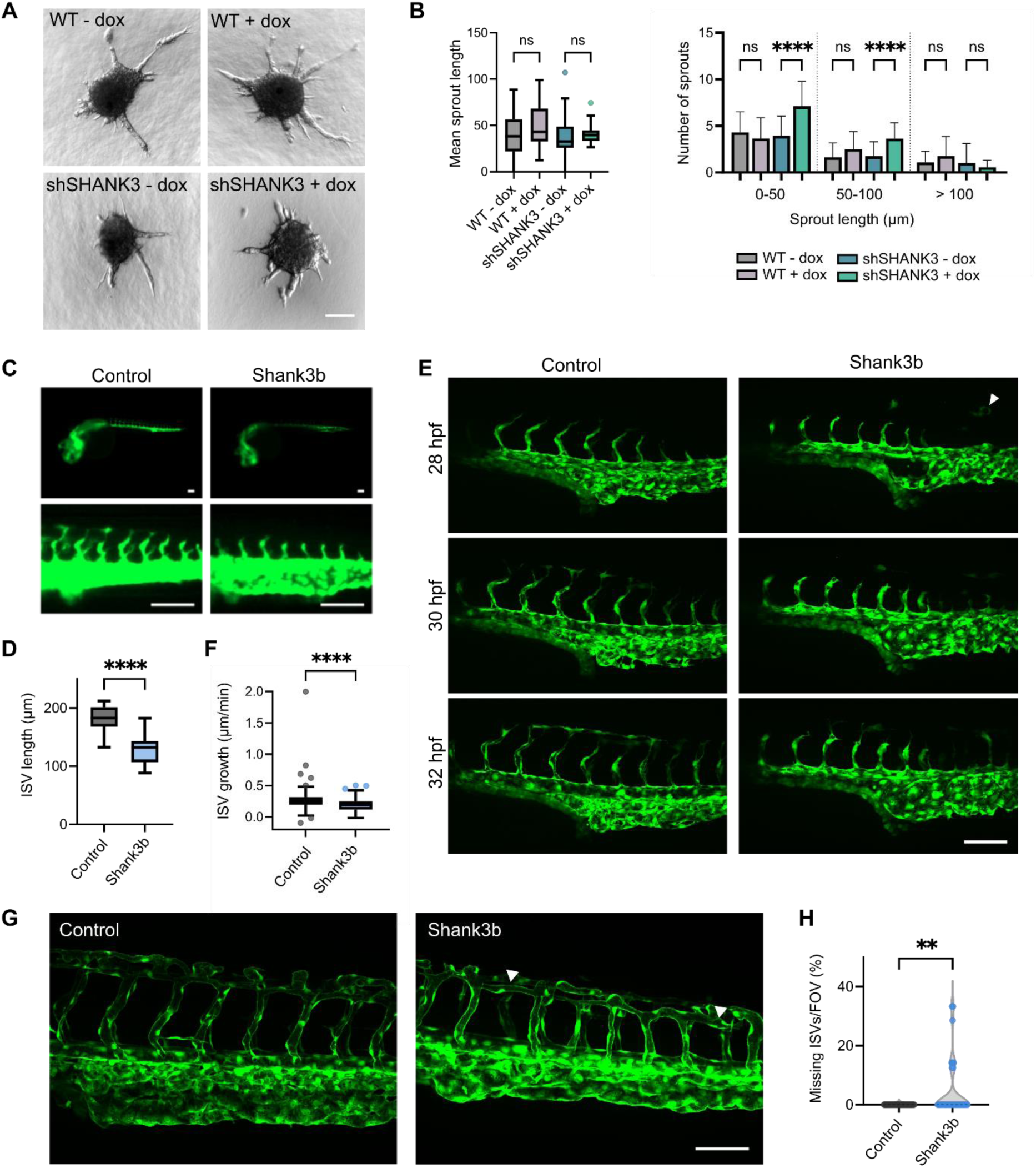
SHANK3 regulates angiogenesis in 3D sprouting and zebrafish embryos. **A.** WT and shSHANK3 HUVEC spheroids +/− doxycycline (dox) were cultured in collagen I gels for 24 h to allow sprouting. Scale bar 100 µm. **B.** Quantification of mean sprout length and number of sprouts (binned by sprout length) per spheroid from A. n = 5 biological replicates, 3-21 spheroids per condition per replicate. One-way ANOVA with Šídák test for multiple comparisons used for statistical analysis. **C.** Fluorescence images of ISVs in control and *shank3b* crispant *Tg(kdrl:EGFP^s843^)* zebrafish at 30 hpf. Endothelial cells shown in green. Scale bar 200 µm. **D**. Quantification of ISV length from C. n = 5 ISVs measured from 24 control and 20 *shank3b* crispants. Unpaired t-test. **E.** Imaging of ISV growth in control and *shank3b* crispant embryos at 26 hpf over 4 h. White arrow indicates abnormally localised endothelial cell. Scale bar 500 µm. **F.** Quantification of ISV growth speed (µm/min) of control and *shank3b* crispant embryos imaged for four hours at 26-32 hpf. n = 2 independent experiments, 57-79 ISVs measured across 10-12 embryos per condition per repeat. Unpaired t-test used for statistical analysis. **G.** Imaging of ISV architecture in control and *shank3b* crispant embryos at 48 hpf. White arrows indicate missing ISVs. Scale bar 500 µm. **H.** Quantification of missing ISVs in G, as percentage of total ISVs per field of view (FOV). n = 3 independent experiments, 8-12 embryos per condition each experiment. Mann-Whitney test used for statistical analysis. ****, p value < 0.001.

Zebrafish have two *shank3* homologues, *shank3a* and *shank3b*. Comparison of their expression with *kdrl* (the zebrafish orthologue of VEGFR2), a marker of endothelial cells, revealed that *shank3b* is highly expressed in endothelial cells (Supplementary Fig. 7A). Consistent with this, *shank3b* expression was detected in endothelial cells from 14-21 hours post fertilisation (hpf) and onwards (Supplementary Fig. 7B).

To explore the role of *shank3b* in the developing zebrafish vasculature, we examined the formation of intersegmental vessels (ISVs) in control or *shank3b* zebrafish crispants (F0 mutant embryos generated with CRISPR-Cas9). ISV formation is an angiogenic process that begins around 22 hpf, during which endothelial cells sprout from the dorsal aorta, migrate dorsally between somites, and ultimately connect to form the dorsal longitudinal anastomotic vessel (DLAV) (Ellertsdóttir et al., 2010). Quantification of the ISV length in control and *shank3b* crispants at 30 hpf revealed a reduced ISV length in *shank3b*-depleted embryos (Fig. 6C and D), indicating a delay in ISV development. Live imaging of embryos between 28 and 36 hpf further revealed reduced migration speed of endothelial cells within developing ISVs in *shank3b* crispants relative to controls (Fig. 6E, F, Supplementary video 2). In addition, in some *shank3b* crispants, atypically localised endothelial cells were also observed in the dorsal region prior to ISV sprouting, disconnected to existing vasculature (Fig. 6E, Supplementary video 2), suggesting potential defects in vascular organisation. Consistent with impaired vascular development, examination of the morphology of ISVs at 48 hpf revealed an increase in the number of missing ISVs in *shank3b* crispants compared to controls (Fig. 6G, H). Together, these results indicate that *shank3b* is important for proper development of the zebrafish vasculature.

We then investigated the role of endothelial Shank3 *in vivo* using the postnatal mouse retina, a well-established model of sprouting angiogenesis. In this system, the primary vascular plexus expands radially from the optic nerve head during the first postnatal week. Whole-mount staining of postnatal mouse retina revealed Shank3 expression in VE-cadherin positive endothelial cells of capillaries and veins, and, to a lesser extent, arteries (Fig. 7A-C, Supplementary Fig. 8A). Furthermore, SHANK3 partially localised to endothelial cell–cell junctions, as indicated by its partial colocalization with VE-cadherin (white arrows, Fig. 7A-C). Endothelial Shank3 deletion was induced by administering 4-hydroxytamoxifen (4-OHT) at postnatal days (P) 1–3 to Shank3^flox/flox^;tdTomato; Cdh5-CreERT2 pups (Shank3^iECKO^) (Fig. 7D). Because CreER^T2^ expression can exert retinal toxicity during development, we compared vascular growth at P6 between 4-OHT-induced Shank3^iECKO^ pups and heterozygous Shank3^Flox/wt^;tdTomato;Cdh5-CreER^T2^ littermate controls (Shank3^iECKO/+^). Tamoxifen-induced tdTomato expression uniformly labelled the vasculature, confirming efficient CreER^T2^ activity in both groups (Fig. 7E). Endothelial-specific deletion of Shank3 in Shank3^iECKO^ retinas was confirmed by immunofluorescence imaging, which showed an almost complete loss of SHANK3 signal in the vasculature, verifying efficient and specific knockout of endothelial Shank3 (Supplementary Fig. 8B).

**Figure 7.**
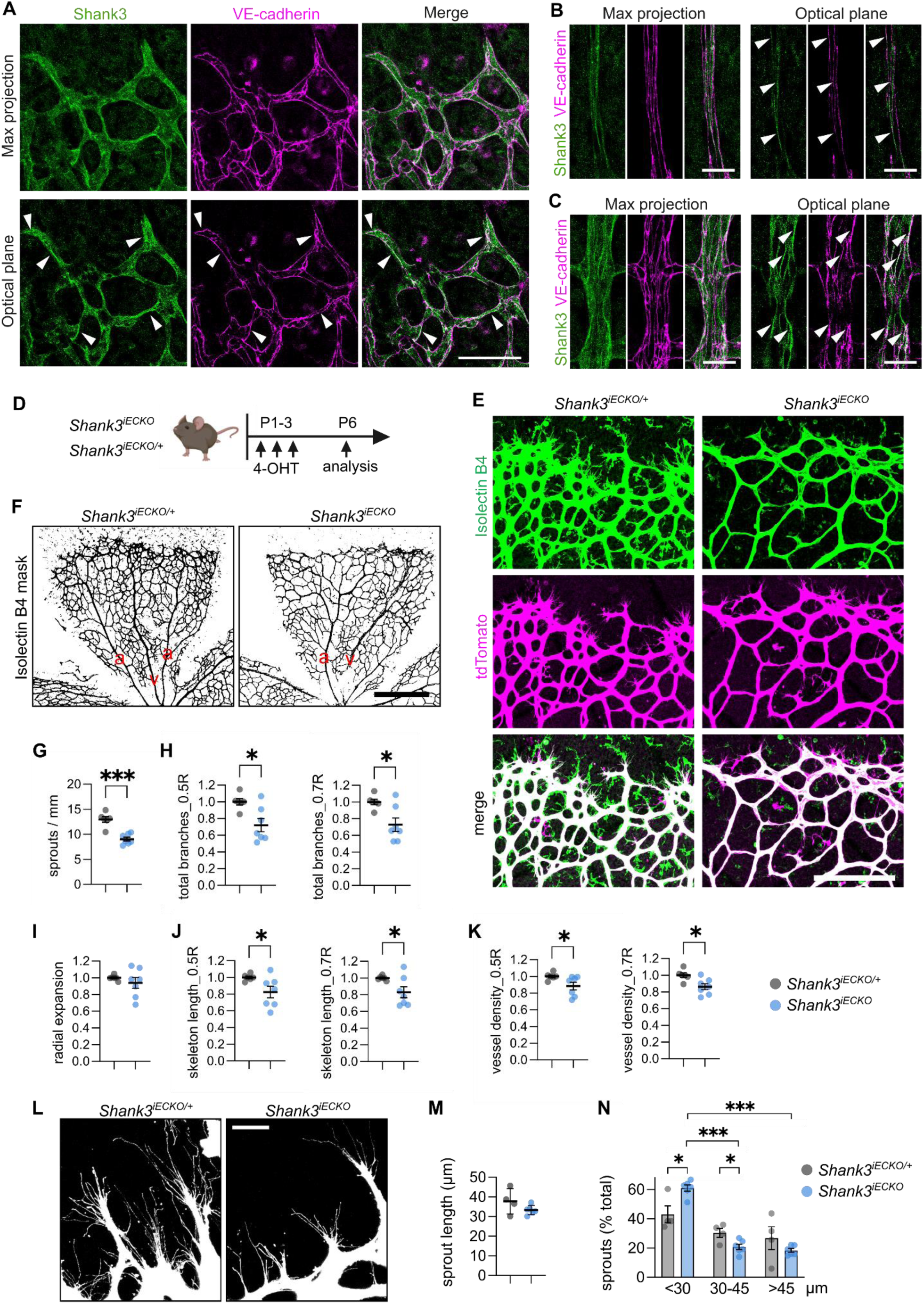
Decreased vascular sprouting in postnatal retina *Shank3^iECKO^*. A-C. Retinal blood vessels (A, vascular front; B, artery; C, vein) from postnatal day 6 (P6) mice stained for Shank3 and VE-cadherin. White arrows indicate colocalization of Shank3 with VE-cadherin at endothelial cell–cell junctions in veins, capillaries, and arteries. Scale bars: 50 µm (A), 20 µm (B, C). **D.** Shank3 deletion was induced using 4-hydroxytamoxifen (4-OHT) in *Shank3*^flox/flox^ (*Shank3*^iECKO^) and *Shank3*^flox/wt^ in tdTomato;Cdh5-CreER^T2^ background (*Shank3*^iECKO/+^) during postnatal days (P) 1–3, followed by analysis of retinal vasculature at P6 (n = 6–7 mice per genotype). Image generated in Biorender.com. **E, F.** Representative images of Isolectin B4 staining and tdTomato expression in retinal vasculature from *Shank3*^iECKO^ and *Shank3*^iECKO/+^ pups. a, artery; v, vein. Scale bars: 200 µm (E), 500 µm (F). **G.** Quantification of sprouts per mm at P6 in the leading front from *Shank3*^iECKO^ and *Shank3*^iECKO/+^. **H-K.** Quantification of the retinal vasculature using the SproutAngio tool (Beter et al., 2023). Shown are total number of branches (H), radial expansion of the vasculature (I), skeleton length (J), and vessel density (K) at r = 0.5R and r = 0.7R, representing proximal and distal zones of the retinal vasculature with respect to optic nerve head (see Supplementary Fig. 8E). Results were normalised to the *Shank3*^iECKO/+^ within litter. **L.** Representative images of sprouts and filopodia expressing tdTomato from *Shank3*^iECKO^ and *Shank3*^iECKO/+^ control pups given 4-OHT at P2–3, followed by analysis of retinal vasculature at P6. Scale bar 20 µm. **M.** Analysis of total sprout length (n = 4–6 mice, described in L). **N.** Sprout length distribution (% of total) showing decreased longer sprouts in *Shank3*^iECKO^ compared to *Shank3*^iECKO/+^ control. n = 4–6 mice (described in L). Statistical tests: two-tailed unpaired t-test (G-K, M), two-way ANOVA with Tukey’s multiple comparison (N). Data are presented as mean ± SEM. P values < 0.05 are shown. ***, p value < 0.005; *, p value < 0.05.

Although radial outgrowth of the vascular plexus was comparable between genotypes (Fig. 7I, Supplementary Fig. 8C-E), Shank3^iECKO^ retinas displayed a clear reduction in sprout number at the angiogenic front, fewer total branch points, shortened vessel skeleton length, and a modest reduction in vessel density (Fig. 7E-K). These alterations were most pronounced at the leading edge (excluding the central retina and optic nerve region), indicating that endothelial Shank3 is specifically required for efficient sprouting and vascular network elaboration, while leaving the overall radial advancement of the vascular front largely intact. Further quantification of sprout length following endothelial-specific deletion of Shank3 revealed a selective loss of longer sprouts (Fig. 7L-N). Although the average sprout length was not significantly altered, *Shank3*^iECKO^ retinas displayed an increased proportion of short sprouts and markedly fewer long sprouts. This pattern mirrored the spheroid sprouting phenotype in 3D gels and suggests that initial sprout formation is largely intact, whereas coordinated cell migration and sprout elongation are impaired in the absence of Shank3. Overall, SHANK3 depletion in both mouse and zebrafish models impairs vascular sprouting during development, indicating an important role in the development of the vasculature.

## Discussion

In this study, we show that SHANK3 is highly expressed in endothelial cells, where it localises to cell-cell junctions. Depletion of SHANK3 leads to altered cell morphology and barrier integrity, and increased cell migration in 2D monolayers. This increase in cell migration in 2D models correlates with altered cell-cell junctions, resulting in reduced tissue viscosity. Interestingly, *in vivo*, SHANK3 depletion led to reduced endothelial cell sprouting and migration, demonstrating that SHANK3 plays an important role in vascular development.

Despite the high expression of SHANK3 in endothelial cells, it is only very recently that its role in the vasculature has begun to be explored. Indeed, SHANK3 was recently shown to be expressed in specialised brain endothelial cells that form the blood brain barrier, where it localised to tight junctions (Y.-E. Kim et al., 2025). The authors demonstrated that SHANK3 depletion contributes towards autism-like behaviour in male neonatal mice through increased permeability of the blood brain barrier, resulting from wnt/β-catenin signalling-mediated disruption of brain endothelial cell tight junctions through reduced expression of tight junction proteins ZO-1 and Claudin5. In our study, we show that SHANK3 is expressed in multiple endothelial cell types across different tissues, suggesting that SHANK3 plays a broader role in endothelial cells beyond the blood brain barrier. Indeed, we show that SHANK3 localises to cell junctions and regulates barrier permeability in HUVECs. Maintaining tight regulation of endothelial cell-cell junctions is essential for the vascular barrier function to control passage of solutes, small molecules and blood cells, and barrier disruption is involved in varied pathologies (Claesson-Welsh et al., 2021). Whether SHANK3 is involved in barrier function in vascular pathologies (beyond contributing to autism-like behaviour in male mice via the blood brain barrier) remains to be seen. Furthermore, SHANK3 has also been shown to regulate barrier function via tight junctions in other cell types. For example, SHANK3 regulates ZO-1 expression in intestinal epithelial cells, with SHANK3 knockout mouse models presenting with an impaired gut barrier function (Wei et al., 2017). This highlights a role for SHANK3 in inflammatory bowel disease and suggests that SHANK3 plays a role in maintaining barriers in multiple tissues, not just in endothelial cells. Further examination into SHANK3 interacting proteins, such as those identified in this study could reveal additional proteins involved in SHANK3 regulation of biological barriers in endothelial cells and other cell types.

Beyond barrier function, we show that SHANK3 depletion alters cell migration in endothelial cells. Cell migration is key to endothelial cell function during development and homeostasis, and is required for vasculogenesis, angiogenesis, and blood vessel repair (Michaelis, 2014). In 2D models, we show that SHANK3 depletion led to increased cell migration in HUVEC monolayers, coupled to an elongated cell shape typically associated with migration (Bi et al., 2015; Blauth et al., 2021; Park et al., 2015; X. Wang et al., 2020). The behaviour of cells in monolayers and the transition between solid- and liquid-like (jammed and unjammed, respectively) has been particularly studied in epithelial cells and tissues. These transitions are important for the development of epithelial tissues, but also diseases such as cancer, where fluid-like states have been linked to cancer dissemination and an invasive phenotype (Blauth et al., 2021; Palamidessi et al., 2019; X. Wang et al., 2020). Cell jamming transitions have been less well studied in endothelial cells. Characterisation of HUVEC migration in monolayers revealed a ‘fluid-like’ way of moving with low intercellular coordination and alignment, rather than coordinated collective migration whereby cell-cell contacts are maintained. Although SHANK3 depletion did not significantly alter the overall number of local cell rearrangements in the monolayer, we observed heterogeneity in migration speeds within the monolayer, with areas of high and low velocity/coordination between cells. This may point towards a transition in the collective behaviour of the monolayer. Collective cell migration involves a complex interplay between cell-ECM adhesion, cell-cell junctions, and contractility (Blauth et al., 2021; Pinheiro & Mitchel, 2024). Altered cell-cell adhesion is known to affect the intracellular tension and tissue mechanics (Pinheiro & Mitchel, 2024). Indeed, in this study, we show that SHANK3 depletion in HUVECs leads to disrupted α-catenin junctions, with reduced levels of α-catenin under high tension. Previously, knockdown of cell-cell components α-catenin and VE-cadherin were shown to increase cell migration in endothelial cell monolayers, without affecting the migration of pioneer cells at leading edges of wounds (Vitorino & Meyer, 2008). This could explain why although increases in migration in HUVEC monolayers are observed upon SHANK3 depletion, scratch-wound healing rates were not affected.

Perhaps initially seemingly in contrast to increased migration in 2D monolayers, we observe reduced endothelial sprouting and migration *in vivo*. However, the modes of migration in 2D monolayers *in vitro* with a flat surface compared to a 3D complex microenvironment with complex topology and chemomechanical signalling cues from the ECM and neighbouring cells are very different (Yamada & Sixt, 2019). *In vivo* and in 3D models, new endothelial cell sprouts depend on the formation of leading tip cells, and their coordination with following stalk cells. Although endothelial sprouting and migration is multifaceted and dependent upon many factors, we hypothesise that the reduced sprouting following SHANK3 depletion may result from defects in cell-cell junctions and intracellular tension between endothelial cells, in addition to reduced traction forces required for pulling on the ECM. Indeed, altered endothelial sprouting and migration have been observed in zebrafish and mice with depletion or disruption of other cell junction components. For example, in zebrafish, knockout of vinculin (which binds α-catenin at VE-cadherin junctions) in endothelial cells resulted in delayed ISV growth and an inability to form junctional fingers (transient, VE-cadherin, ZO-1 and PECAM-1-positive structures that form at cell-cell junctions)(Kotini et al., 2022), while VE-cadherin has been shown to be required for dynamic stalk cell behaviour (Sauteur et al., 2014). In mice, deletion of vinculin or β-catenin in endothelial cells also led to reduced vessel density and sprouting (Carvalho et al., 2019; Martowicz et al., 2019). Interestingly, an increased prevalence of vision and eye disorders have been identified in people with ASD, in which mutations in SHANK3 is a known factor (Perna et al., 2023). Some of the eye disorders are associated with altered vasculature in the retina, including reduced peripheral vision and reduced contrast sensitivity (Ameri et al., 2025; Liu et al., 2022). Although it is not yet known whether these particular cases are due to altered vasculature or whether they are linked to SHANK3 mutations, it would be interesting to further investigate the possible link between vascular development in the retina and SHANK3 in ASD.

Overall, we demonstrate that SHANK3 plays an important function in endothelial cells, affecting barrier formation, cell migration and vascular development, with implications for vascular disease, in addition to shankopathies such as ASD.

## Methods

### scRNAseq biofinformatics

The Tabula Sapiens single-cell RNA-sequencing dataset and associated metadata were obtained from the Gene Expression Omnibus (GSE201333). Preprocessed, normalised gene expression matrices and scVI-harmonised UMAP embeddings were used for downstream analyses and visualization (Jones et al., 2022). Predefined tissue, compartment, and cell ontology annotations were used as grouping variables. All analyses and visualizations were performed in R using the Seurat and ggplot2 packages.

### Staining of FFPE tissue sections

H&E-stained sections were obtained from routine surplus formalin-fixed paraffin-embedded (FFPE) diagnostic control tissue, irreversibly anonymised as part of the diagnostic workflow. The material was not required for clinical use and the sectioned slides routinely discarded after a defined expiration period of 30 days following sectioning.

Staining was performed using the Sakura Tissue-Tek Prisma system. Briefly, sections were deparaffinised in xylene (2 × 3 min) and rehydrated through a graded ethanol series (100%, 96%, and 70%; 2 min each), followed by rinsing in running tap water and immersion in distilled water (1 min each). Hematoxylin staining was performed using Tissue-Tek Hematoxylin 3G (Sakura) for 5 min. After washing in running water (1 min) and differentiation in 70% ethanol (30 s), sections were stained with Tissue-Tek Eosin (Sakura) for 1.5 min. Finally, sections were dehydrated through an ascending ethanol series (70%, 96%, and 100% for 1, 2, and 3 min, respectively) and cleared in xylene (2 min followed by 3 min).

### Cell lines and culture

Human Umbilical Vein Endothelial Cells (HUVEC; PromoCell, pooled donors) were cultured in Endothelial Cell Growth Medium (ECGM) with SupplementMix (PromoCell, C-22010) and 1% penicillin-streptomycin (Sigma). HUVECs were not used beyond 4 passages. Human osteosarcoma U2OS cells (DSMZ, ACC 785) were cultured in Dulbecco’s Modified Eagle Medium (DMEM) supplemented with 10% fetal bovine serum (FBS, Biowest) with 1% penicillin-streptomycin and 1% L-glutamine (Sigma). Cells were maintained at 37°C in a humidified atmosphere with 5% CO_2_.

### siRNAs, plasmids and cloning

For SHANK3 depletion by siRNA, two individual Human SHANK3 siRNAs were used: siRNA#2 (Hs_SHANK3_2 siRNA, Qiagen, target sequence 5’-CAGGGATGTCCGCAACTACAA-3’) and siRNA#7 (ON-TARGETplus^TM^ Hs_SHANK3_7 siRNA, Dharmacon^TM^ Horizon Discovery, target sequence: 5’-GGGCTTCACCTGACTACAA-3’). AllStars negative control siRNA (Qiagen) was used as a non-targeting siRNA control. pPB-TagBFP-T2A-myc-BirA*-SHANK3 (BirA-SHANK3) and pPB-TagBFP-T2A-SHANK3-myc-BirA* (SHANK3-BirA) BioID constructs were generated using NEBuilder HiFi DNA Assembly (NEB). For BirA-SHANK3, SHANK3 was amplified from GFP-SHANK3 (source) using forward (GAAGCGCAGAGAAGCTCGAGAGACCTCTAATGGACGGCCCCGG) and reverse (CACAGTGGCGGCCGCTCGAGTCAGCTGCCATCCAGCTGCC) primers, and inserted into pPB-TagBFP-T2A-myc-BirA* (pPB-BirA) (ref eplin paper) linearised with XhoI. For SHANK3-BirA, SHANK3 was amplified from GFP-SHANK3 (source) using forward (AGAATCCCGGCCCTTCCGGAATGCAGCTAAACCGTGCCGCCG) and reverse primers (CTGAGATGAGTTTTTGTTCTGAGCCGCCTCCTCCAGCTTTGCTGCCATCCAGCT), and inserted into pPB-BirA linearised with BspEI. pTriEx-mCherry-Zdk1 and pTriEx-TOM20-Ven-Lov2 were a gift from Klaus Hahn. Shank3-mCherry-Zdk1-N3 was generated by GenScript by inserting mCherry-Zdk1 into Shank3-mRFP-N3 plasmid.

### Generation of cell lines

SMARTvector^TM^ shRNA lentiviral particles (shSHANK3, Cat. no. V3SH7669-228381856, target sequence ATACAAGCGGCGAGTTTAT, Dharmacon) were used to generate doxycycline-inducible shRNA SHANK3-depleted HUVECs (more info in (Lilja et al., 2024)). The shSHANK3 lentiviral vector contains a TRE3G inducible promotor activated by Tet-ON 3G transactivator protein in the presence of doxycycline, inducing expression of the shRNA and a turboRFP reporter (allowing visual tracking of shRNA expression). For stable expression of shSHANK3 in HUVECs, p0 HUVECs were incubated for 24 h with ECGM (without pen/strep) containing packaged lentivirus (30 MOI) and polybrene (8 µg/mL, Sigma). Three days following transduction, cells were selected with 1 µg/ml puromycin (Sigma) for 48 h. For shSHANK3 induction, cells were incubated with 1 µg/ml doxycycline 6 h after seeding, and culture media (+/− dox) was changed daily. Cells were used for experiments three days following initial doxycycline addition.

Stably-expressing BioID U2OS cell lines (pPB-BirA, pPB-BirA-SHANK3, and pPB-SHANK3-BirA) were generated by co-transfection of pPB-BioID constructs with pCMV-HypBase (a plasmid containing the piggyBac transposase (yusa et al., 2018)) at a 3:1 ratio (pPB-BioID vector:pCMV-HypBase) with Lipofectamine 3000 according to the manufacturers protocol. Culture media was replaced with fresh media the following day. Cells were sorted for high and low BFP expression two weeks later using FACS, and BirA-protein expression confirmed using western blotting and subcellular localisation confirmed using immunofluorescence imaging. Although highly expressing BirA control cells were selected, BirA expression levels were significantly lower than that of BirA-SHANK3 and SHANK3-BirA expressing cells.

### siRNA silencing

HUVEC cells were reverse transfected with 30 nM siRNA#2, siRNA#7 or AllStars negative control using Lipofectamine RNAiMAX (ThermoFisher) according to manufacturer’s instructions. For transfection in 6-well plates, 5 x 10^5^ cells in 500 µl ECGM was added to 500 µl siRNA/Lipofectamine RNAiMAX mix in Opti-MEM (ThermoFisher) and incubated for 2 h at 37°C before an additional 500 µl ECGM added to the wells. 24 h after transfection, media was changed to 2 ml full culture medium. For transfection in 24-well and 96-well plates, transfection reagents and cell numbers were scaled accordingly. Transfected cells were used for experiments at indicated time points.

### Immunofluorescence imaging of fixed cells

Cells were fixed with 4% paraformaldehyde (PFA) in PBS for 15 min at RT before washed 3 times with PBS. Cells were permeabilised with 0.2% Triton-X-100 in PBS for 5 mins before PFA quenched with 0.1M glycine for 30 min. For the FN-accessibility assay to examine barrier function, cells were not permeabilised (Ball et al., 2024). Cells were incubated with primary antibodies (see table 2 for details) in 10% horse serum in PBS overnight at 4°C, washed three times with PBS, and incubated with Alexa Fluor-conjugated secondary antibodies (Alexa Fluor 488-, 568- and 647-conjugated anti-mouse, anti-rabbit and anti-rat, 1:500, Life Technologies), and/or Phalloidin-Atto 647 (1:500, Sigma) for 1 h at RT. Cells were then washed three times with PBS and incubated with 4’,6-diamidino-2-pheylindole (DAPI, 1:10000 in PBS) for 5 min, and finally washed a further three times with PBS before mounting (Prolong Gold Antifade, Invitrogen) onto glass slides. Images were captured using a Marianas CSU-W1 Spinning disk spinning disk confocal microscope (3i/Zeiss) using a LD C-APOCHROMAT 40x/1.1 W M27 objective (Zeiss) and sCMOS Orca Flash 4.0 camera (Hamamatsu).

### BioID and affinity purification of proteins

pPB-BioID U2OS cells were seeded onto two 10 cm dishes, and the following day biotin (50 µM) added to initiate biotinylation of proximal proteins for 24 h. Cells were then lysed and biotinylated proteins affinity-purified as previously described (Chastney, Lawless, & Humphries, 2020; Chastney, Lawless, Humphries, et al., 2020). Cells were washed three times with 5 ml PBS and cells lysed using 400 µl lysis buffer (250 mM NaCl, 50 mM Tris HCl pH 7.4, 0.5 mM DTT, 0.1 % SDS (w/v), and 1X protease inhibitors (cOmplete Mini, EDTA-free, Roche)). 80 µl 20% Triton-X 100 was added to lysates (800 µl per condition) and passed four times through a 19G needle. Next, 720 µl 50 mM Tris HCl (pH 7.4) was added to lysates, before being passed through a 27G needle four times, and centrifuged 10 min at 16,000 xg at 4°C in a microcentrifuge. The supernatant was rotated overnight at 4°C with 30 µl MagReSyn magnetic streptavidin beads (ReSyn Biosicences). Beads were then washed twice with 500 µl wash buffer 1 (10% SDS (w/v)), once with wash buffer 2 (500 mM NaCl, 50 mM HEPES, 1 mM EDTA, 1% Triton X-100 (w/v), 0.1% deoxycholic acid (w/v)) and once with wash buffer 3 (10 mM Tris HCl (pH 7.4), 1 mM EDTA, 0.5 % NP-40 (w/v), 0.5% deoxycholic acid (w/v)) before biotinylated proteins eluted at 70°C for 10 min in 90 µl 2 X reducing sample buffer with 10 µM biotin.

### Mass spectrometry sample preparation and data acquisition

For in-gel tryptic digestion of BioID samples, eluted proteins were ran through SDS-PAGE until all sample had entered the gel (200V for ∼4 minutes, 2 x wells per condition, 4-20% Mini-PROTEAN TGX protein gel, Bio-Rad). Gels were incubated with Coomassie Blue for ten minutes before being washed with 3 X 10 min washes with ddH_2_O, and a final wash with ddH_2_O overnight at 4°C. Protein bands were excised, cut into 1-2mm pieces, and washed twice with 0.04 M NH_4_HCO_3_/50% acetonitrile (ACN) for 15 min. Gel pieces were then incubated with 100% ACN for 10 min, and proteins reduced for 30 min at 56°C using 20 mM DL-Dithiothreitol (DTT; BioUltra, Sigma). 100% ACN was added to the gel pieces for 10 min, followed by a 20 min incubation at RT in 55 mM iodoacetamide in 100 mM NH_4_HCO_3_ for protein alkylation. Gel pieces were washed twice with 100 mM NH_4_HCO_3_ and once with 100% ACN before centrifugation using a vacuum centrifuge. Proteins were then digested overnight at 37°C using 5 ng/µl sequencing grade modified trypsin (Promega) in 40 mM NH_4_HCO_3_ and 20% ACN. To extract peptides, gel pieces were incubated for 15 min at 37°C in 100% ACN followed by 15 min at 37°C in 50 % ACN/5% formic acid, with supernatant collected after each incubation. Finally, peptides were dried in a vacuum centrifuge. For MS analysis, samples were dissolved in 10 µl 2% formic acid immediately prior to use.

The LC-ESI-MS/MS analyses were performed on a nanoflow HPLC system (Easy-nLC1200, Thermo Fisher Scientific) coupled to the Q Exactive HF mass spectrometer (Thermo Fisher Scientific, Bremen, Germany) equipped with a nano-electrospray ionization source. Peptides were first loaded on a trapping column and subsequently separated inline on a 15 cm C18 column (75 μm x 15 cm, ReproSil-Pur 3 μm 120 Å C18-AQ, Dr. Maisch HPLC GmbH, Ammerbuch-Entringen, Germany). The mobile phase consisted of water with 0.1% formic acid (solvent A) and acetonitrile/water (80:20 (v/v)) with 0.1% formic acid (solvent B). A 30 min linear gradient from 6% to 39% of solvent B, followed by a wash stage with 100% of eluent B was used to eluate peptides. MS data was acquired automatically by using Thermo Xcalibur 4.1 software (Thermo Fisher Scientific). A data dependent acquisition method consisted of repeated cycles of one MS1 scan covering a range of m/z 350 – 1750 plus a series of HCD fragment ion scans (MS2 scans) for up to 10 most intense peptide ions from the MS1 scan.

### MS data processing and bioinformatics

BioID MS data were analysed as previously described (ref). MaxQaunt (v2.0.3.0, available from Max Planck Institute of Biochemistry (Tyanova et al., 2016)(Tyanova et al., 2016)(Tyanova et al., 2016)(Tyanova et al., 2016)) was used for processing Raw MS data, with default parameters and biotinylation of lysines as a valuable modification, LFQ quantification, and match between runs selected. MS/MS spectra were searched against the human proteome obtained from Swiss-Prot (April 2021). The mass spectrometry proteomics data have been deposited to the ProteomeXchange Consortium (http://proteomecentral.proteomexchange.org) via the PRIDE partner repository (Perez-Riverol et al., 2025) with the dataset identifier (to be specified). To identify high-confidence interactors, LFQ intensities from MaxQuant were analysed with SAINTexpress (Teo et al., 2014) using default parameters. Proximity interactors were identified using a BFDR threshold of ≤ 0.05. ClusterProfiler (version 4.8.3) was used to perform Gene ontology analysis (Yu et al., 2012). Interactome networks were identified using STRING and imported into Cytoscape (version 3.10.1) (Su et al., 2014), and keywords identified from UniProt Annotated Keywords. Known SHANK3 interactors were identified from BioGRID (release 4.4.205) (Stark et al., 2006).

### ‘Knock-sideways’ targeting of SHANK3 to mitochondria using LOVTRAP

For examination of mis-localised proteins, U2OS cells were co-transfected with pTriex-TOM20-Ven-Lov-WT and pTriEx-mCherry-Zdk1 (H. Wang et al., 2016) or Shank3-mCherry-Zdk1-N3 (Shank3-mCherry-Zdk1) using Lipofectamine 3000 following manufacturers protocol. The following day, cells were plated onto Ibidi µ-Slide 8-well, fixed 24 h later, and prepared for imaging as described above.

### Western blotting

For cell lysis, cells were maintained on ice and washed twice with ice cold PBS before being lysed with RIPA buffer (50 mM Tris HCl pH 7.4, 150 mM NaCl, 1% Triton X-100, 0.1% sodium deoxycholate, 0.1% SDS, 1 mM EDTA, 10 mM NaF) with 1X cOmplete protease inhibitor cocktail (Sigma) and 1X PhosSTOP phosphatase inhibitors (Sigma). Samples were scraped, transferred to an Eppendorf tube, and sonicated for 10 mins (30 sec on, 30 sec off). Lysates were then centrifuged at 16,000 xg for 20 minutes at 4°C in a tabletop microcentrifuge. Protein concentrations were analysed using a DC Protein assay (Bio-Rad) according to manufacturer’s protocol. Reducing sample buffer was added, and samples boiled for 5 min at 95°C. Proteins were subjected to SDS-PAGE under denaturing conditions (4-20% Mini-PROTEAN TGX gels, Bio-Rad) before being transferred to nitrocellulose membranes using the Trans-Blot Turbo Transfer System (Bio-Rad). Membranes were blocked with AdvanBlock-Fluor blocking solution (Advansta) diluted 1:1 in TBST (Tris-buffered saline with 0.1% Tween20) for 1 h. Membranes were incubated overnight at 4°C with primary antibodies in AdvanBlock-Fluor:TBST blocking solution. Membranes were washed three times with TBST for 5 min before incubation for 2 h at RT with secondary antibodies (1:5000 ECL HRP-conjugated secondary antibodies for pERK; 1:2500 fluorophore-conjugated Azure secondary antibodies, AH Diagnostics, for all other antibodies). Membranes were washed three times with TBST for 5 minutes. Membranes were imaged with an infrared imaging system (Azure Sapphire RGBNIR Biomolecular Imager, Azure Biosystems) or using ECL plus Western blotting substrate (Pierce) and Chemidoc (BioRad).

### RT-qPCR

RNA was extracted from cells using a NucleoSpin RNA kit (Macherey-Nagel), and 1 µg RNA used as a template for cDNA synthesis using a High-Capacity cDNA Reverse Transcription kit (Thermo Scientific). SHANK3 expression levels were examined using Taqman qRT-PCR, using TaqMan® Fast Advanced Master Mix (Applied Biosystems). TaqMan® Gene Expression Assays (Thermo Scientific) were used to detect Shank3 (Hs00873185_m1) and GAPDH (Hs02786624_g1). Results were analysed with QuantStudio. Relative quantification (RQ) of SHANK3 was determined from three technical replicates using GAPDH as an endogenous control.

### Live cell imaging and cell migration tracking

Cells were plated onto 24-well plates, and shSHANK3 induction induced as described above. 3 days following initial doxycycline addition, media was changed to ECGM containing 20 mM HEPES (pH 7.5, Sigma) and spy-650-DNA (1:5000, Spirochrome). Cells were maintained at 37°C and 5% CO_2_, and imaged every 10 minutes for 16 h using a Nikon Eclipse Ti2-E with a Plan Fluor 10x objective (Nikon) and an Orca Flash 4.0 sCMOS camera (Hamamatsu Photonics). For scratch wound assays, a clean pipette tip was used to remove cells.

For cell tracking, masks of cell nuclei were first generated using a custom trained StarDist 2D model using ZeroCostDL4Mic (Schmidt et al., 2018; von Chamier et al., 2021). The StarDist 2D model was trained for 100 epochs on 11 paired image patches (image dimensions: (2044, 2048), patch size: (2016,2016)) with a batch size of 2 and a mae loss function, using the StarDist 2D ZeroCostDL4Mic notebook (v1.19.1) (von Chamier et al., 2021). The StarDist ‘Versatile fluorescent nuclei’ model was used as a training starting point (Schmidt et al., 2018). Key python packages used include tensorflow (v 2.14.0), csbdeep (v 0.7.4), cuda (v 11.8.89 Build cuda_11.8.r11.8/compiler.31833905_0). The training was accelerated using a Tesla T4 GPU. This approach produced a model demonstrating an average F1-score of 0.959 on our test dataset. Labels were then tracked using the TrackMate plugin in Fiji (Ershov et al., 2022; Schindelin et al., 2012; Tinevez et al., 2017), using the TrackMate label detector and the Advanced Kalman Tracker with a maximum frame gap of 3, and filtering for track duration above 38 time points.

### Migration analysis

The instantaneous velocity 𝒗_𝑖_(𝑡) of each cell was obtained from nuclear tracking as the ratio between the individual cell displacement between two consecutive frames (𝒙_𝑖_(𝑡 + Δ𝑡) − 𝒙_𝑖_(𝑡)) and the time delay Δ𝑡 between frames. Two-point velocity correlation function was calculated as

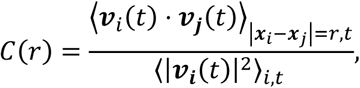

where ⟨⋅⟩_|𝒙_𝑖_−𝒙_𝑗_|=𝑟,𝑡_ indicates an average performed over all time points 𝑡 and all cell pairs at mutual distance 𝑟, and ⟨⋅⟩_𝑖,𝑡_ an average over all time points 𝑡 and all cells. Two-point rms velocity correlation function was calculated as

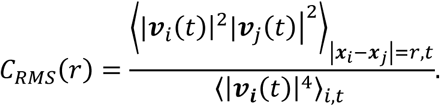

A correlation length 𝐿_𝐶_ was estimated by fitting the model 𝑓(𝑟) = 𝐴 cos(𝜋𝑟/𝐿_𝑐_)𝑒^−𝜋𝑟/𝐿𝑐^ + 𝐵 to 𝐶_𝑅𝑀𝑆_(𝑟).

The frequency 𝑓_𝑇1_ of T1 (neighbour switching) events was estimated as follows. For each time t, we considered the cells whose nuclei are correctly tracked in both frames collected at times 𝑡 and 𝑡 + Δ𝑡, respectively. For each frame, we calculated the Delaunay triangulation of the nuclear centres. This enabled identifying, for each cell, its nearest neighbours, as the cells connected to it by an edge of the triangulation. The number 𝑁(𝑡) of T1 events occurred between frames 𝑡 and 𝑡 + Δ𝑡 is obtained as the number of changes occurred in all lists of nearest neighbours the divided by 4 (each T1 event involves 4 cells). Finally, the frequency 𝑓_𝑇1_ was obtained as ⟨𝑁(𝑡)⟩_𝑡_/Δ𝑡.

### PIV analysis of MEK inhibited cells

HUVECs were plated into fibronectin-coated (10 µg/ml) wells of an E-Plate VIEW 96 plate (Agilent Technologies) and transfected with control and SHANK3 targeting siRNAs as described above. Triplicate wells were used. Three days following siRNA knockdown, media was replaced with ECGM +/− 250 nM Trametinib MEK inhibitor (MedChemExpress), and cells imaged every 20 minutes for 24 hours using a xCELLigence RTCA eSight (Agilent Technologies). PIV analysis of monolayers at 23-24 h following MEK inhibition was performed using OpenPIV-Python (Liberzon et al., 2021) via ‘PIV_4_xCelligence’ Jupyter notebook (https://github.com/CellMigrationLab/PIV_4_xCELLigence), and average velocity shown.

### Tubule formation assay

Three wells of Ibidi µ-Plate 96 well per condition were coated with 10 µl Matrigel (Corning), and allowed to polymerise at 37°C for 1 h, before 15,000 cells were plated per well. Cells were imaged 18-20 h after seeding using a Leica Thunder widefield microscope, with 5x N Plan objective and a Leica K8 camera. Images were analysed using the Angiogenesis Analyzer for ImageJ (Carpentier et al., 2020).

### Traction force microscopy

For hydrogels (∼10 kPa), 35 mm glass bottom dishes (D35-14-1, CellVis) were incubated with 1 ml bind silane solution (7.14 % Plus One Bind silane (Sigma) and 7.14 % acetic acid in EtOH) for 30 min at room temperature before being washed twice with EtOH and dried. 1.7 μl sonicated fluorescent beads (200 nm Yellow-Green FluoSpheres^TM^ Carboxylate-Modified Microspheres) were added to 500 μl hydrogel mix, containing 94 μl 40 % acrylamide (Sigma) and 50 μl N,N’-methylenebisacrylamide (Sigma) in PBS, and 5 μl 10 % ammonium persulfate (Bio-Rad) and 1 μl N,N,N’N’-tetramethylethylenediamine (Sigma) was added to the hydrogel mixture to induce polymerisation. The mixture was vortexed and 11.8 μl added to the dishes, and a 13 mm glass coverslip placed on top. After a 1 h incubation at room temperature, PBS was added to the dishes and the glass coverslips removed. The hydrogel surface was activated by 30 min incubation at room temperature in 500 μl 0.2 mg/ml Sulfo-SANPAH (Sigma) in 50 mM HEPES, followed by 10 min irradiation with UV light. Gels were washed four times with PBS and coated with 10 μg/ml fibronectin for 2 h at room temperature.

Cells were seeded onto hydrogels 24 h, and 2 h prior to imaging media was replaced with media +/−doxycycline containing 20 mM HEPES (pH 7.0-7.6, Sigma), 2 µg/ml Hoescht (Invitrogen), and 200 nM SiR-actin (Spirochrome) to enable visualisation of cells. For detection of traction forces exerted by the cells, beads were imaged before and after removal of cells with 15 µl pre-warmed 20% SDS, with a Marianas CSU-W1 Spinning disk spinning disk confocal microscope (3i/Zeiss) using a LD C-APOCHROMAT 40x/1.1 W M27 objective (Zeiss) and sCMOS Orca Flash 4.0 camera (Hamamatsu). To correct for drift, ‘before’ and ‘after’ images were aligned using the Fast4Dreg plugin in Fiji (Pylvänäinen et al., 2023; Schindelin et al., 2012). Tracking of beads and force measurements were analysed using TFM software (Han et al., 2015) in MATLAB (Mathworks, version 2024a). Grid points and PIV suite was used for displacement field calculation, with a template size of 21 pixels and maximum displacement of 20 pixels. For displacement field correction, vector outliers were filtered and a threshold of 2 for the normalised displacement residual. Force field calculation was performed using Fourier transform traction cytometry (FTTC) and a regularisation parameter of 0.0001.

### Spheroids and associated assays

To generate spheroids, 3000 HUVECs were seeded into wells of ultra-low attachment round bottomed 96 well plates (Corning) in 200 µl media +/− 1 µg/ml doxycycline and incubated for 24 h. For spheroid wetting assays, spheroids were re-seeded into 24-well glass-bottomed plates coated with 10 µg/ml fibronectin in media containing 20 mM HEPES (pH 7.5) +/− 1 µg/ml doxycycline, and imaged every 10 min for 12 h at 37°C and 5% CO_2_ using a Nikon Eclipse Ti2-E with a Plan Fluor 10x objective (Nikon) and an Orca Flash 4.0 sCMOS camera (Hamamatsu Photonics). For spheroid merge assays, two spheroids were transferred to a single well of an ultra-low attachment round bottomed 96 well plate, and media replaced with fresh ECGM with 20 mM HEPES and +/− 1 µg/ml doxycycline. Spheroids were maintained at 37°C and 5% CO_2_ and imaged every 10 min for 12 h using a Leica Thunder widefield microscope with a 10x HC Plan Apo objective and Leica K8 camera. For spheroid sprouting assays, 3-5 spheroids were seeded into 50 µl rat tail collagen I (Corning; final concentration 2 mg/ml) containing 1X DMEM and NaOH added to induce polymerisation. Collagen was allowed to polymerise at 37°C for 1 h before 150 µl media +/− 1 µg/ml doxycycline added. Spheroids were incubated at 37°C and 5% CO_2_ for 24 h before imaging with a Leica Thunder widefield microscope with a 10x HC Plan Apo objective and Leica K8 camera.

### Zebrafish crispants

crRNAs targeting *shank3b* were designed using ChopChop (Labun et al., 2019). *Shank3b* crRNAs and Alt-R™ Cas9 Negative Control crRNA #1 were ordered from Integrated DNA Technologies (IDT). Control or *shank3b* crRNAs and Alt-R™ CRISPR-Cas9 tracrRNA (IDT) at equimolar concentrations were combined in nuclease-free duplex buffer (IDT) to final concentrations of 25 µM using annealing program of 5 min at 95°C, cooling 0.1°C/sec to 25°C, 5 min at 25°C and 4°C on hold. Injection mix consisting of 5 µM duplex, 0.8 µg/µl Alt-R™ S.p. Cas9 Nuclease V3 (IDT), 240 mM KCl and 10% phenol red in nuclease-free water was prepared fresh before injection and incubated 5 min at 37°C. The zebrafish (strain *Tg(kdrl:EGFP^s843^)* obtained through timed natural spawning in specific mating tanks were collected with a plastic sieve, rinsed with fish system water and transferred to a petri dish. The 1-4 cell stage embryos were injected with 2.3 nl injection solution into the yolk using a Nanoject II injector (3-000-205A, Drummond). Embryos were incubated in 0.0025% 1-phenyl 2-thiourea (PTU) and 50 U/ml Penicillin-Streptomycin in E3 (5 mM NaCl, 0.17 mM KCl, 0.33 mM CaCl2, 0.33 mM MgSO4) at 28.5°C. Successful editing was confirmed by PCR followed by Sanger sequencing and chromatogram analysis using TIDE (Brinkman et al., 2014).

### Zebrafish imaging and quantification

For measurement of ISV length, widefield imaging of zebrafish ISVs was performed using a Nikon Ti-2e inverted microscope equipped with a Hamamatsu Orca Flash 4.0 sCMOS camera. Images were acquired using a Plan Apo λ 4× objective (NA 0.2). Multi-channel imaging was performed using GFP emission filter (emission 515/30 nm) and fluorescence excitation was provided by a Lumencor Spectra light source using 475 nm (50% power) excitation line. Vessel length was measured manually from dorsal edge of dorsal aorta to the distal tip of the migrating ISV using manual measurement in ImageJ 1.54f/FIJI (Schindelin et al., 2012). For ISV migration, 26-32 hpf embryos were mounted (as described previously (Follain et al., 2018)) and ISVs imaged every 10 min for 4 h at 28.5°C using a Marianas CSU-W1 Spinning disk spinning disk confocal microscope (3i/Zeiss) with a 25x/0.8 Imm Korr DIC objective (Zeiss) and a sCMOS Orca Flash 4.0 camera (Hamamatsu). ISVs were tracked until branching/connecting to the DLAV. For quantification of missing ISVs, 48 hpf embryos were mounted and imaged taken from the post-cloacal region using Marianas CSU-W1 Spinning disk spinning disk confocal microscope (3i/Zeiss) with a 25x/0.8 Imm Korr DIC objective (Zeiss) and a sCMOS Orca Flash 4.0 camera (Hamamatsu).

### Mouse models and preparation of whole-mount retina

This study examined both male and female mice, and the findings were similar for both sexes. The following mouse lines were used: *Shank3^flox/flox^* (B6.Cg-Shank3tm1.1Bux/J, Jackson Laboratory, stock no 017889) (Bozdagi et al., 2010), *Rosa26-tdTomato^lox/STOP/lox^* (The Jackson Laboratory) and the previously published *Cdh5-CreER^T2^* (Tg(Cdh5-cre/ERT2)Ykub) (Okabe et al., 2014) all maintained in pure C57BL/6 background (>10 backcrosses). Gene deletions were induced by administration of 4-hydroxytamoxifen (4-OHT; Sigma, H6278) dissolved in ethanol and corn oil (i.p. 20 ug g^-1^ x 3 days) at P1-P3 or P2-P3 and analysed at P6. *Shank3^flox/flox^;Cdh5-CreER^T2^*pups (*Shank3^iECKO^*) were compared to 4-OHT-treated *Cre*-positive *Shank3^Flox/+^;Cdh5-CreER^T2^* controls, to control for potential Cre-ER^T2^ mediated toxicity, as reported previously (Brash et al., 2020). Number of mice per experiment are indicated in figures. For whole-mount staining of postnatal mouse retina, eyes were fixed in 4% PFA for 1 h or 2 h at +4°C. Prior to staining, four radial incisions were made in the retina to allow flat-mounting. The tissues were blocked in donkey immunomix (DIM; 5% donkey serum, 1% BSA, 0.3% Triton X-100, 0.05% sodium azide in Dulbecco’s PBS), incubated with primary antibodies in DIM for 1-3 days at 4°C, washed with 0.3% TritonX-100 in PBS, incubated with secondary antibodies O/N at RT and washed. Biotinylated isolectin B4 (IB4) staining was performed in Pblec buffer (1 mM MgCl_2_, 1 mM CaCl_2_, 0.1 mM MnCl_2,_ 1% TritonX-100 in PBS) O/N at +4°C, followed by washing with 0.3% TritonX-100 in PBS, incubation with streptavidin-conjugated Alexa Fluor dyes in DIM O/N at RT, and washing with 0.3% TritonX-100 in PBS. Whole mounts were post-fixed with 4% PFA for 10 min, washed with PBS and mounted with DAPI containing Vectashield mounting medium (H-1200, Vector Labs). Laser scanning confocal Z-stack images were acquired using Leica Stellaris 8 Falcon confocal microscope (HC PL APO 10x /NA 0.4 objective, 63x /NA 1.40 oil objective or 40x /NA 1.25 glycerol objective) or Andor Dragonfly 505 spinning disc microscope (Plan Apo 20x /NA 0.75 objective). All confocal images represent maximum intensity projections of Z-stacks if not indicated otherwise.

### Analysis of mouse retina using SproutAngio

Automated quantification of retinal vasculature was performed using the SproutAngio tool(Beter et al., 2023), which segments Isolectin B4–labelled vessels and extracts structural parameters from the skeletonized vascular network. Images were processed with uniform thresholding and segmentation settings across samples. For each retina, the central region containing the optic nerve was excluded to isolate the angiogenic front, following SproutAngio’s radial-mask-based workflow. The tool computes branch points from the skeleton graph, radial expansion from the maximum migration distance of vessels, and total skeleton length from the cumulative length of all vessel segments. Vessel density was calculated as the proportion of vascular area relative to the total tissue area within the defined radial zones (r = 0.5R and r = 0.7R), obtained by dividing the binarized vessel area by the area of the corresponding region of interest. All metrics were normalized to *Shank3*^iECKO/+^ littermate controls to reduce inter-litter variability. The length and number of sprouts at the angiogenic front was quantified manually from 20x tile images or 63x images and normalized to the length of the vascular front.

## Acknowledgements

We thank Jenni Siivonen and Petra Laasola for excellent technical assistance, and the members of the Ivaska laboratory, especially Johanna Lilja, for valuable scientific discussion. Imaging was performed at the Advanced Imaging Core Facility at Turku Bioscience Centre, supported by Biocentre Finland, the Finnish Advanced Microscopy Node of Euro-BioImaging Finland (Turku, Finland), and Turku Bioimaging, and was supported by the Research Council of Finland, FIRI 2023 grant decision numbers 359073 and 358879, and FIRI 2024 grant decision numbers 367582 and 367577. Testament funds from Henna Ruusunen also supported this work. We also thank the Turku Zebrafish Core, the Turku Proteomics Facility and the Turku Cytometry Core at Turku Bioscience Centre, all supported by Biocentre Finland. This work was supported by an ERC Advanced Grant (BorderControl; grant No. 101142305 to J.I.). This work has been supported by the Finnish Cancer Institute (K. Albin Johansson Professorship, J.I. and G.J.), a Research Council of Finland Centre of Excellence (grant numbers 346131 and 364182, J.I. and P.S.), the Cancer Foundation Finland (J.I.), the Sigrid Juselius Foundation (J.I. and G.J.), the Research Council of Finland’s Flagship InFLAMES (grant numbers 337530, 357910, 337531, and 35791), the Jane and Aatos Erkko Foundation (J.I.), the Research Council of Finland (grant numbers 343239 to M.R.C., 338537, 371287, and 374180 to G.J., and 332402 to G.F.), the Cancer Society of Finland (Syöpäjärjestöt; G.J.), the Solutions for Health strategic funding for Åbo Akademi University (G.J.), and the Turku Collegium for Science, Medicine, and Technologies (G.F.). Funded by the European Union. Views and opinions expressed are however those of the author(s) only and do not necessarily reflect those of the European Union or the European Research Council Executive Agency. Neither the European Union nor the granting authority can be held responsible for them.

## Author contributions

Conceptualisation, M.R.C and J.I.; methodology, M.R.C., A.P., J.H., J.W.P., F.G., I.P.; formal analysis, M.R.C., A.P., J.H., V.S., F.G., I.P.; investigation; M.R.C., A.P., J.H., G.F., V.S., M.V., A.M.H.S., J.V., I.P.; writing – original draft, M.R.C. and J.I.; writing – review and editing, M.R.C., A.P., J.H., G.F., V.S., J.W.P., A.M.H.S., J.V., M.V., G.S., I.P., G.J., F.G., P.S., and J.I.; supervision, G.S., G.J., F.G., P.S. J.I.; funding acquisition, M.R.C., J.I.

