## Supplemental Data for "Scaffold protein SHANK3 regulates endothelial cell motility and tissue mechanics"

**Supplementary Figures and Figure legends:**

Supplementary Figure 1, Related to Figure 1.

Supplementary Figure 2, Related to Figure 1.

Supplementary Figure 3, Related to Figure 2.

Supplementary Figure 4, Related to Figure 3.

Supplementary Figure 5, Related to Figure 4.

Supplementary Figure 6, Related to Figure 5.

Supplementary Figure 7, Related to Figure 6.

Supplementary Figure 8, Related to Figure 7.

**Supplementary tables**

Supplementary Table 1. SHANK3 proximity interactome

Supplementary Table 2. List of antibodies used

**Supplementary Videos:**

Video 1: Live imaging of HUVECs labelled with SiR-Actin and GFP-CAAX showing ‘aster’-like actin structures.

Video 2: Live imaging of control and *Shank3b*-depleted zebrafish embryos showing development of vasculature between 28-32 hpf.

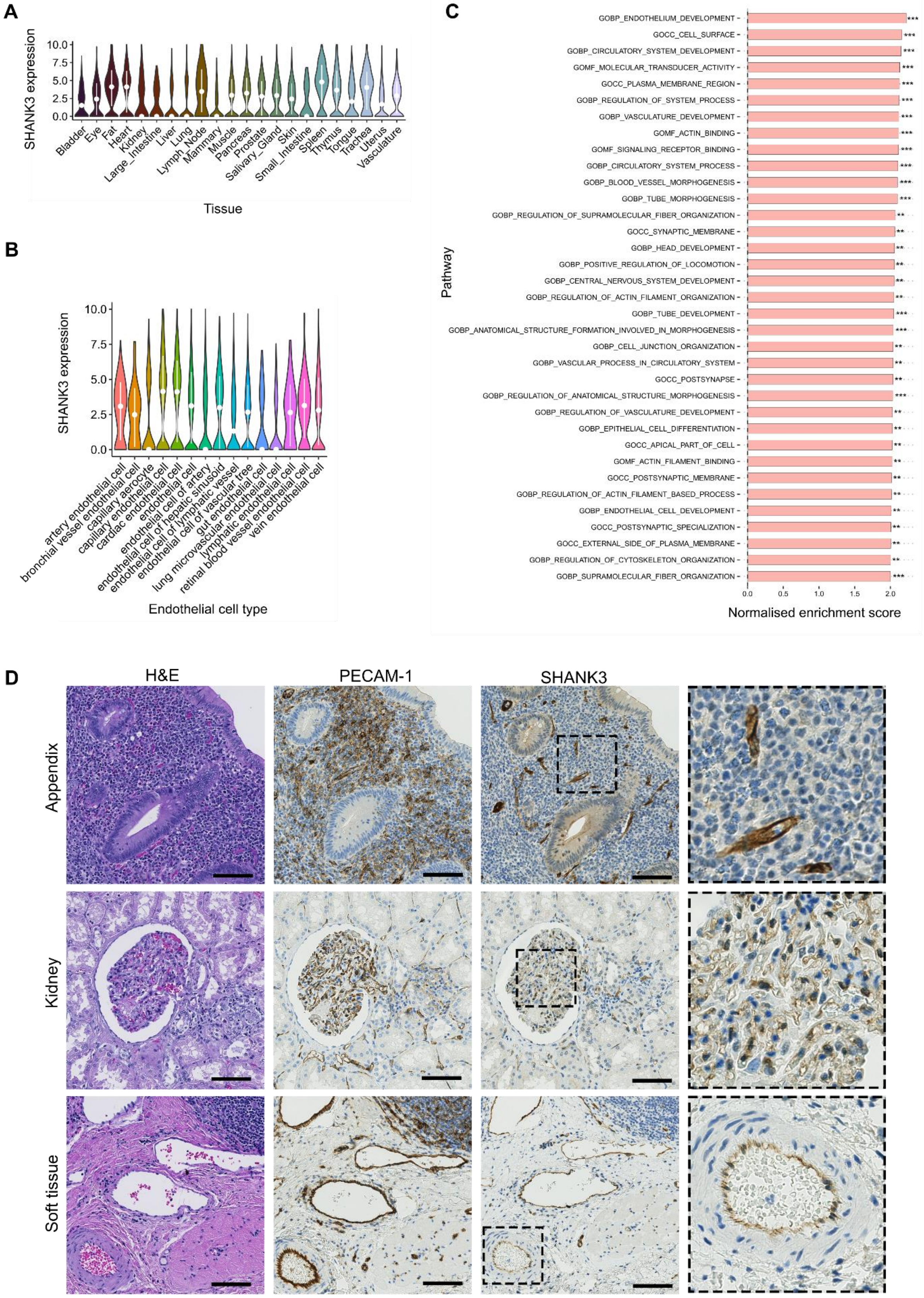
A
SHANK3 expression
10.0
7.5
5.0
2.5
0.0
Bladder
Eye
Fat
Heart
Kidney
Large_Intestine
Liver
Lung
Lymph_Node
Mammary
Muscle
Pancreas
Prostate
Salivary_Gland
Skin
Small_Intestine
Spleen
Thymus
Tongue
Trachea
Uterus
Vasculature
Tissue
B
SHANK3 expression
10.0
7.5
5.0
2.5
0.0
artery endothelial cell
bronchial vessel endothelial cell
capillary aerocyte
capillary endothelial cell
cardiac endothelial cell
endothelial cell
endothelial cell of artery
endothelial cell of hepatic sinusoid
endothelial cell of lymphatic vessel
endothelial cell of vascular tree
gut endothelial cell
lung microvascular endothelial cell
lymphatic endothelial cell
retinal blood vessel endothelial cell
vein endothelial cell
Endothelial cell type
C
Pathway
GOBP_ENDOTHELIUM_DEVELOPMENT
GOCC_CELL_SURFACE
GOBP_CIRCULATORY_SYSTEM_DEVELOPMENT
GOMF_MOLECULAR_TRANSDUCER_ACTIVITY
GOCC_PLASMA_MEMBRANE_REGION
GOBP_REGULATION_OF_SYSTEM_PROCESS
GOBP_VASCULATURE_DEVELOPMENT
GOMF_ACTIN_BINDING
GOMF_SIGNALING_RECEPTOR_BINDING
GOBP_CIRCULATORY_SYSTEM_PROCESS
GOBP_BLOOD_VESSEL_MORPHOGENESIS
GOBP_TUBE_MORPHOGENESIS
GOBP_REGULATION_OF_SUPRAMOLECULAR_FIBER_ORGANIZATION
GOCC_SYNAPTIC_MEMBRANE
GOBP_HEAD_DEVELOPMENT
GOBP_POSITIVE_REGULATION_OF_LOCOMOTION
GOBP_CENTRAL_NERVOUS_SYSTEM_DEVELOPMENT
GOBP_REGULATION_OF_ACTIN_FILAMENT_ORGANIZATION
GOBP_TUBE_DEVELOPMENT
GOBP_ANATOMICAL_STRUCTURE_FORMATION_INVOLVED_IN_MORPHOGENESIS
GOBP_CELL_JUNCTION_ORGANIZATION
GOBP_VASCULAR_PROCESS_IN_CIRCULATORY_SYSTEM
GOCC_POSTSYNAPSE
GOBP_REGULATION_OF_ANATOMICAL_STRUCTURE_MORPHOGENESIS
GOBP_REGULATION_OF_VASCULATURE_DEVELOPMENT
GOBP_EPITHELIAL_CELL_DIFFERENTIATION
GOCC_APICAL_PART_OF_CELL
GOMF_ACTIN_FILAMENT_BINDING
GOCC_POSTSYNAPTIC_MEMBRANE
GOBP_REGULATION_OF_ACTIN_FILAMENT_BASED_PROCESS
GOBP_ENDOTHELIAL_CELL_DEVELOPMENT
GOCC_POSTSYNAPTIC_SPECIALIZATION
GOCC_EXTERNAL_SIDE_OF_PLASMA_MEMBRANE
GOBP_REGULATION_OF_CYTOSKELETON_ORGANIZATION
GOBP_SUPRAMOLECULAR_FIBER_ORGANIZATION
0.0
0.5
1.0
1.5
2.0
Normalised enrichment score
D
H&E
PECAM-1
SHANK3
Appendix
Kidney
Soft tissue

**Supplementary figure 1. SHANK3 is expressed in multiple endothelial cell subtypes**

**A.** Expression of SHANK3 mRNA in endothelial cells of multiple tissues. Mean ± s.d. shown in white. **B.** SHANK3 mRNA expression in endothelial cell subtypes. Mean ± s.d. shown in white. **C.** GSEA pathway enrichment in SHANK3 mRNA expressing endothelial cells relative to non-expressing endothelial cells. ***, p value < 0.001; ** p value < 0.005. Data from the Tabula Sapiens (Jones et al., 2022). **D.** Histological staining of serial sections of appendix, kidney, and soft tissue. PECAM-1 used as endothelial cell marker. Scale bar 100 µm. H&E, hematoxylin and eosin.

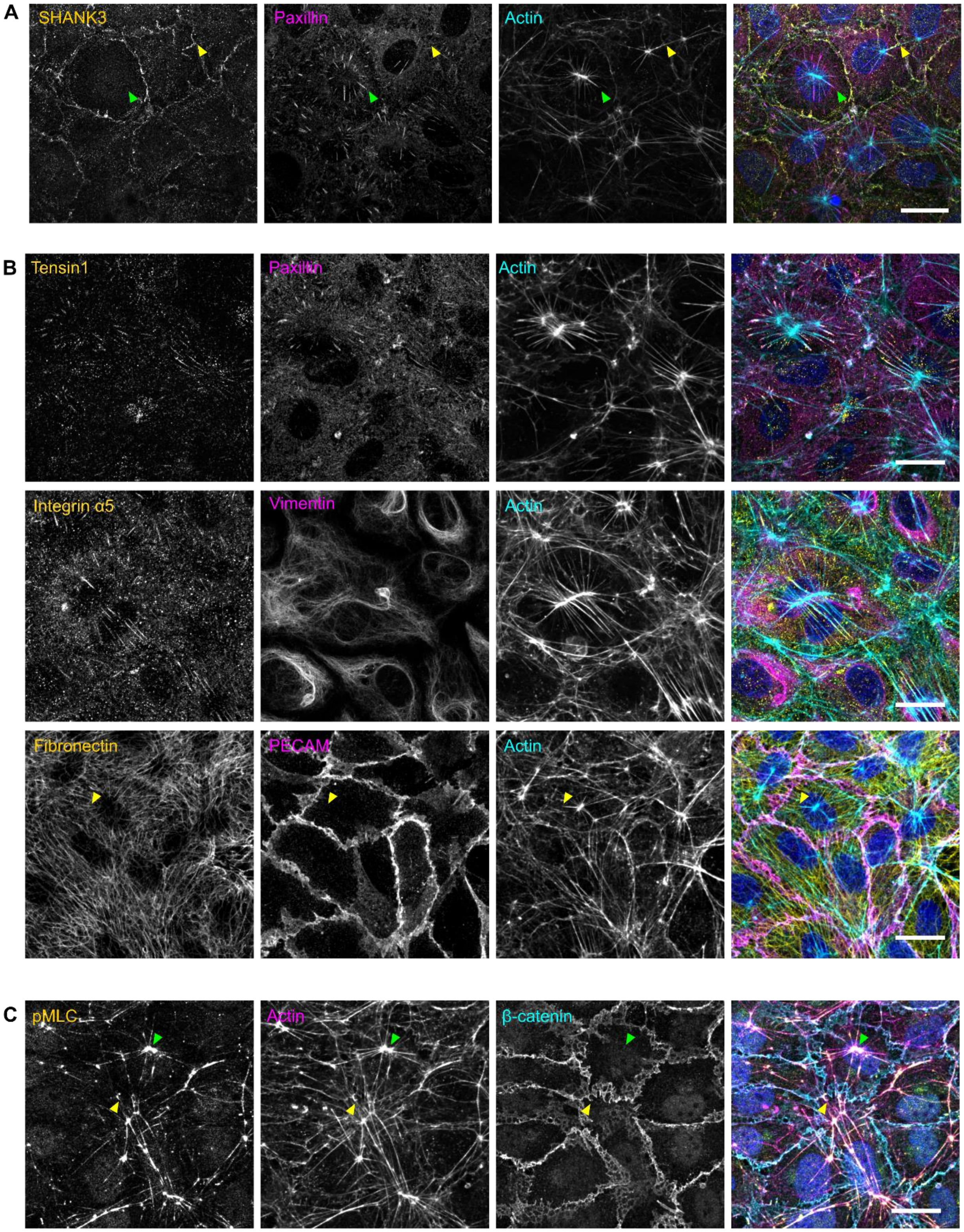
A
SHANK3
Paxillin
Actin
B
Tensin1
Paxillin
Actin
Integrin α5
Vimentin
Actin
Fibronectin
PECAM
Actin
C
pMLC
Actin
β-catenin

**Supplementary figure 2. SHANK3 associates with contractile actin structures linking cell-cell and cell-ECM contacts**

**A.** Immunofluorescence imaging of HUVEC monolayers show 'aster/spoke'-like actin structures that connect to both paxillin-positive integrin adhesion complexes (green arrows) and SHANK3-positive cell-cell junctions that link actin between two cells (yellow arrows). Scale bar 20 µm. **B.** Immunofluorescence imaging of HUVEC monolayers stained with fibrillar adhesion markers tensin-1 and integrin α5 (SNAKA51 antibody) and fibronectin. Yellow arrows indicate fibronectin fibres continuous with aster-like actin structures. Paxillin used as a general marker for integrin adhesion complexes, PECAM-1 shows cell-junctions, and vimentin staining also shown. Scale bar 20 µm. **C.** phospho-Myosin Light Chain (pMLC) staining at the centre of aster-like actin structures (green arrows) and at actin at cell junctions (yellow arrows) suggest that these structures are highly contractile. Scale bar 20 µm.

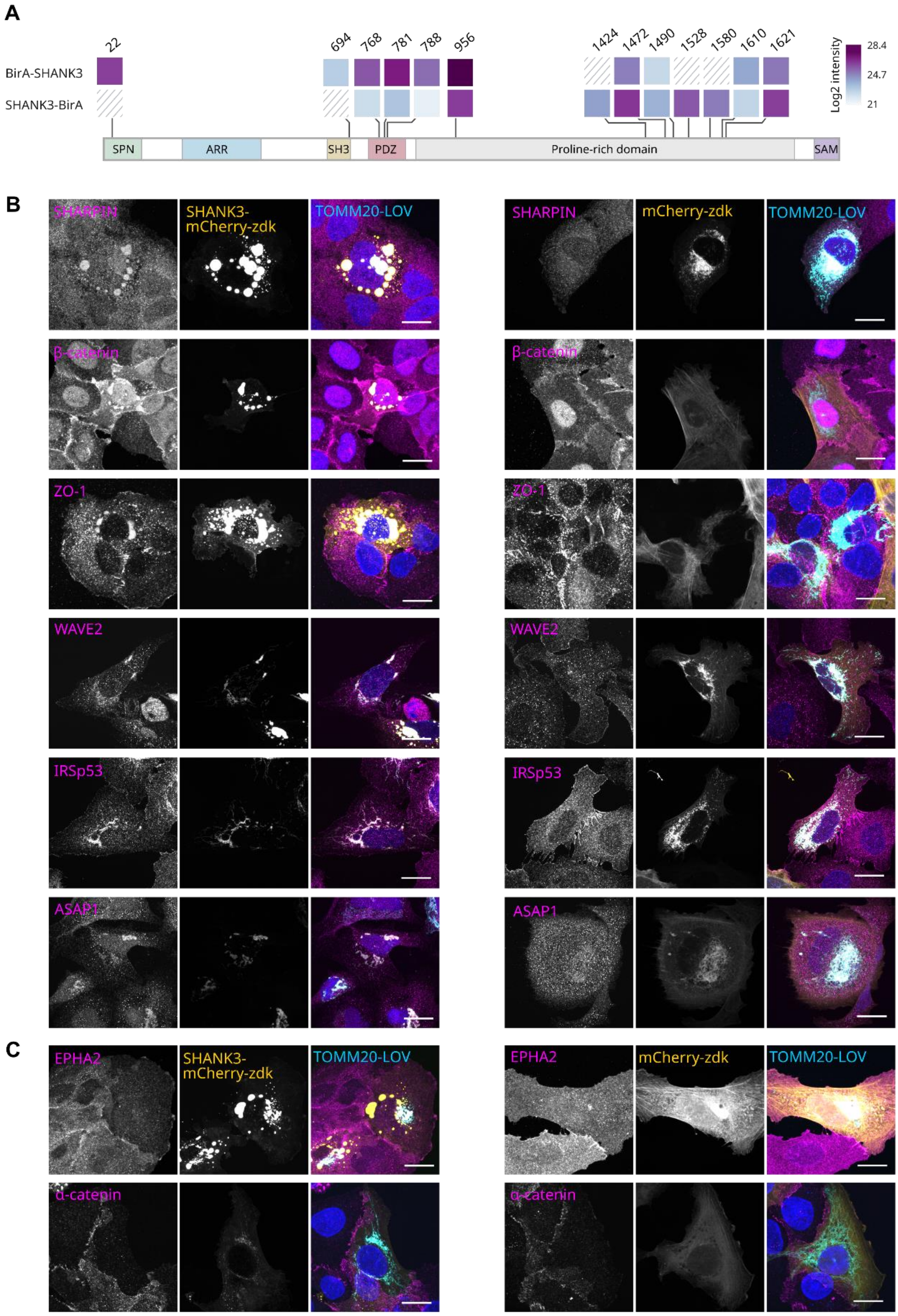

A
BirA-SHANK3
SHANK3-BirA
22
694
768
781
788
956
1424
1472
1490
1528
1580
1610
1621
Log2 intensity
28.4
24.7
21
SPN
ARR
SH3
PDZ
Proline-rich domain
SAM
B
SHARPIN
SHANK3-mCherry-zdk
TOMM20-LOV
mCherry-zdk
β-catenin
ZO-1
WAVE2
IRSp53
ASAP1
C
EPHA2
α-catenin

**Supplementary figure 3. SHANK3 biotinylated peptides and validation of proximity interactors**

**A.** Biotinylated peptides in SHANK3 identified by BirA-SHANK3 and SHANK3-BirA. Biotinylated site locations are shown above and mapped onto the SHANK3 domain structure shown below. SPN, Shank/ProSAP N-terminal domain; ARR, Ankyrin Repeat Region; SH3, Src Homology 3 domain; PDZ, PSD95, DlgA and ZO-1 domain; SAM, Sterile Alpha Motif. **B.** Immunofluorescence imaging showing mis-localisation of SHANK3 interactors to mitochondria following a knock-sideways approach. SHANK3-mcherry-zdk is targeted to mitochondria via TOMM20-LOV, and SHANK3 interactor candidates stained using antibodies. mCherry-zdk used as a negative control. Scale bar 20 µm. **C.** Immunofluorescence images of SHANK3 interactor candidates showing no mis-localisation to mitochondria via SHANK3-mCherry-zdk and TOMM20-LOV. Scale bar 20 µm.

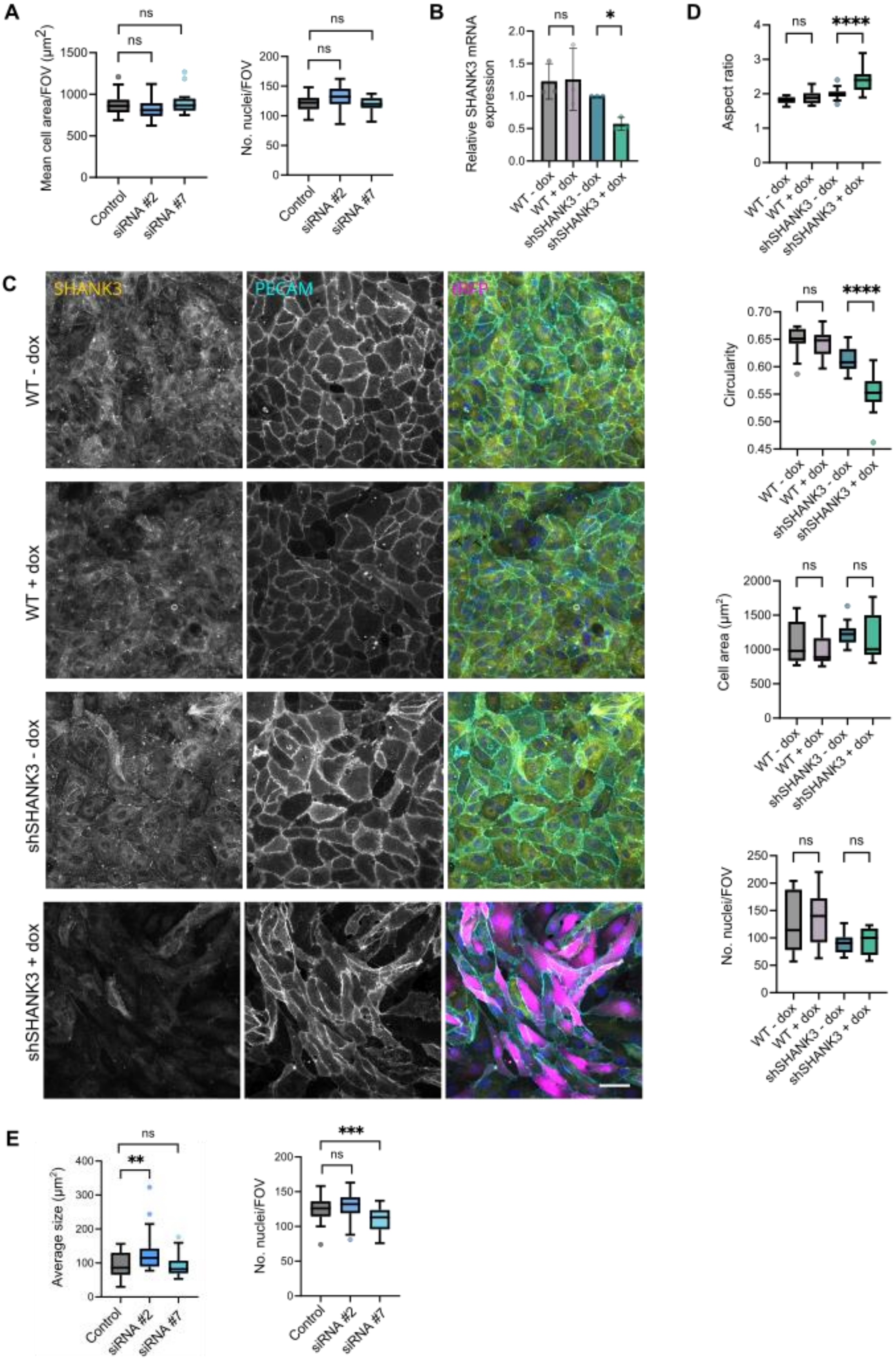

A
B
C
D
E
ns
*
****
***
**
Mean cell area/FOV (μm²)
No. nuclei/FOV
Relative SHANK3 mRNA expression
Aspect ratio
Circularity
Cell area (μm²)
Average size (μm²)
Control
siRNA #2
siRNA #7
WT - dox
WT + dox
shSHANK3 - dox
shSHANK3 + dox
SHANK3
PECAM
RFP

**Supplementary figure 4. SHANK3 depletion using shRNA leads to altered cell morphology**

**A.** Quantification of mean cell circularity and number of nuclei per field of view from Fig. 3B. n = 3 biological replicates, 6 fields of view per condition each replicate. Unpaired t-test. **B.** Relative SHANK3 mRNA expression in WT HUVECs and HUVECs expressing doxycycline-inducible anti-SHANK3 shRNA (shSHANK3) +/- doxycycline (dox). Expression normalised to shSHANK3 - dox samples. n = 3 biological replicates. One-way ANOVA with Šídák test for multiple comparisons used for statistical analysis. **C.** Immunofluorescence imaging of WT and shSHANK3 HUVECs +/- doxycycline. tRFP signal indicates expression of shRNA construct. Scale bar 20 µm. **D.** Quantification of cell shape (PECAM-1) and number of nuclei per field of view from B. n = 3 biological replicates, 5-7 fields of view per condition each replicate. One-way ANOVA with Šídák test for multiple comparisons used for statistical analysis. **E.** Quantification of fibronectin patch size and number of nuclei per field of view in Fig. 3D. Unpaired t-test. n = 3 biological replicates, 10-13 fields of view per condition each replicate. ns, non-significant; ****, p value < 0.01; *, p value < 0.05.

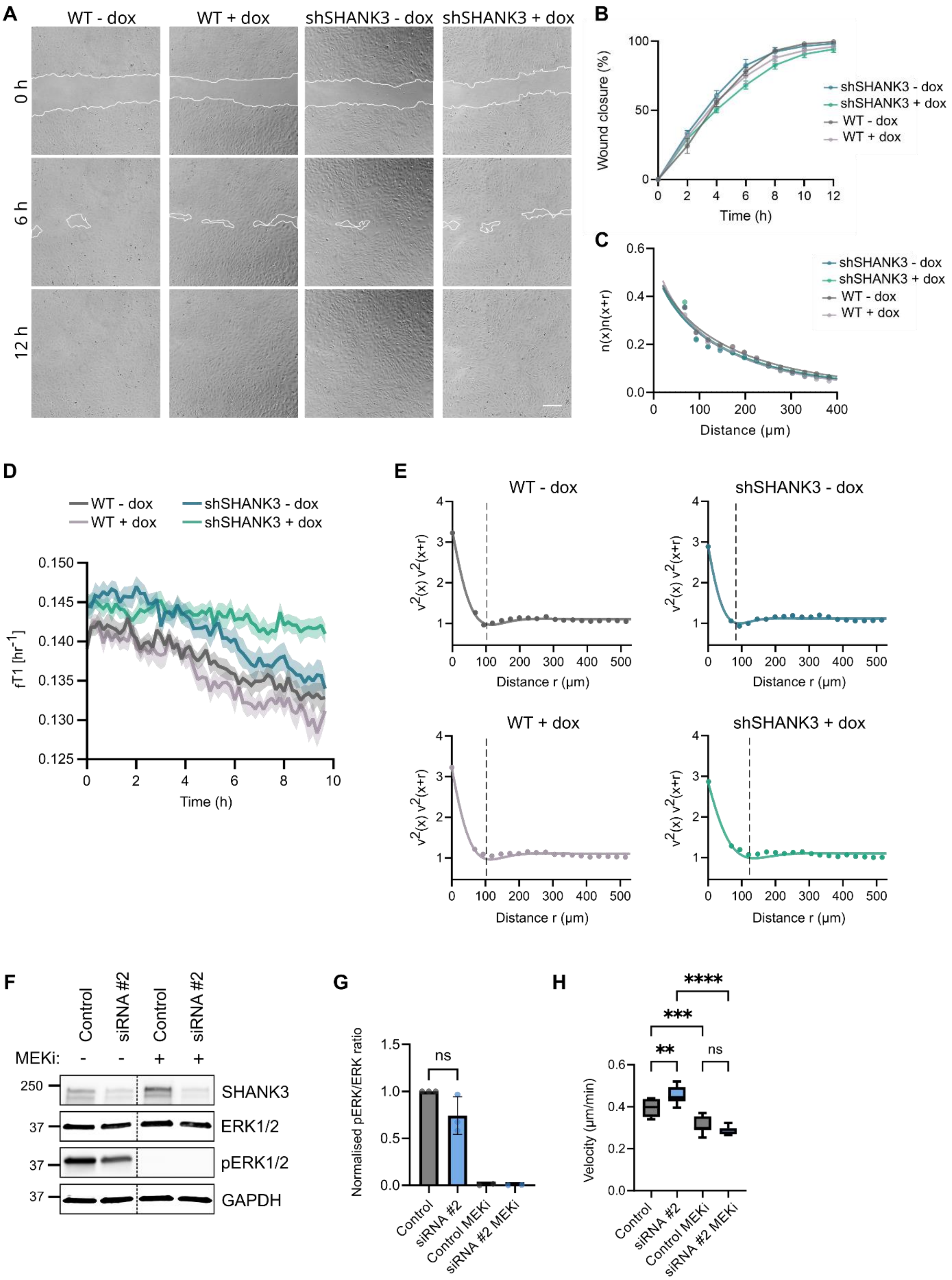

A
WT - dox
WT + dox
shSHANK3 - dox
shSHANK3 + dox
0 h
6 h
12 h
B
Wound closure (%)
Time (h)
shSHANK3 - dox
shSHANK3 + dox
WT - dox
WT + dox
C
n(x)n(x+r)
Distance (μm)
D
fT1 [hr-1]
Time (h)
E
v2(x) v2(x+r)
Distance r (μm)
F
Control
siRNA #2
MEKi:
SHANK3
ERK1/2
pERK1/2
GAPDH
G
Normalised pERK/ERK ratio
ns
Control MEKi
siRNA #2 MEKi
H
Velocity (μm/min)
****
***
**
ns

**Supplementary figure 5. Effects of ERK and Rap1 inhibition on cell migration in SHANK3-depleted HUVECs**

**A.** Scratch wound assay showing wound closure of WT and shSHANK3 HUVECs +/- doxycycline (dox) over 12 h. Scale bar 200 µm. **B.** Quantification of wound closure in A. One way ANOVA with Šídák test for multiple comparisons used for statistical analysis at each time point (all non-significant), mean ± SEM shown. n = 3 biological replicates, 3-6 fields of view per condition each replicate. **C.** Two-point velocity correlation function $\langle \boldsymbol{n}(\boldsymbol{x}) \cdot \boldsymbol{n}(\boldsymbol{x}+\boldsymbol{r}) \rangle$ from WT and shSHANK3 HUVECs +/- dox. Mean of 5 fields of view from one representative replicate shown. **D.** Mean frequency of T1 events of cells ($f_{T1}$) per field of view. Mean ± SEM shown. n = 3 biological replicates, 5 fields of view per condition in each replicate. **E.** Two-point correlations in the root mean square velocity $\langle v^2(\boldsymbol{x}) v^2(\boldsymbol{x}+\boldsymbol{r}) \rangle$. Dashed line indicates the estimated correlation length. Mean of 5 fields of view from one representative replicate shown. **F.** Western blot showing levels of ERK phosphorylation following MEK inhibition (trametinib, MEKi) in WT and SHANK3 siRNA-depleted HUVECs. Dotted line indicates cropping of blot. **G.** Quantification of pERK levels from A, normalised to ERK. n = 3 for untreated samples, 2 for MEKi treated samples. Unpaired t-test used for statistical analysis of untreated samples. **H.** Velocity of WT and SHANK3 siRNA-depleted HUVECs following ERK pathway inhibition with MEKi. Control, AllStars negative control; ns, non-significant; ****, p value < 0.001; ***, p value < 0.005; **, p value < 0.01.

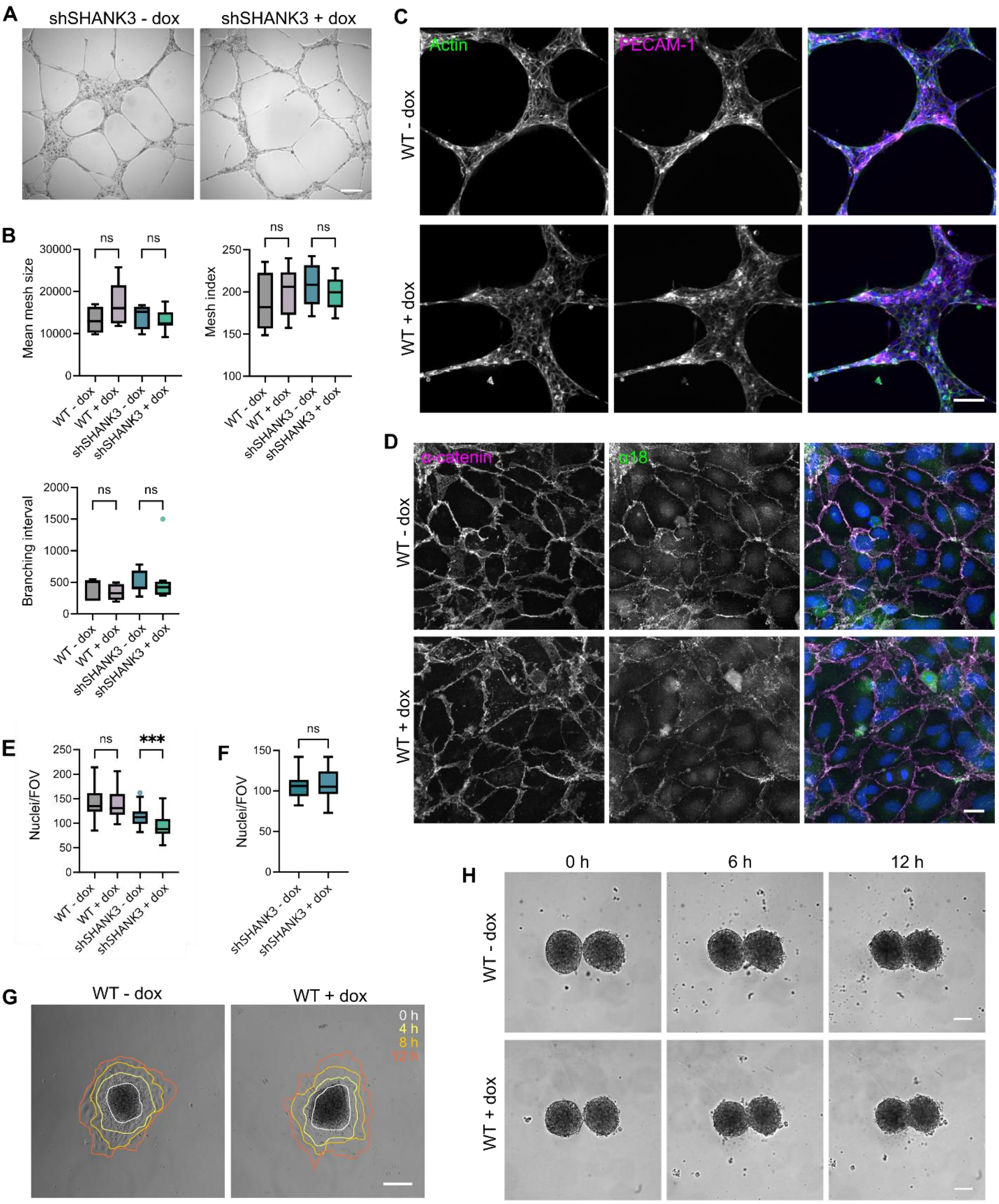
A
shSHANK3 - dox
shSHANK3 + dox
B
Mean mesh size
30000
20000
10000
0
ns
ns
WT - dox
WT + dox
shSHANK3 - dox
shSHANK3 + dox
Mesh index
250
200
150
100
Branching interval
2000
1500
1000
500
C
Actin
PECAM-1
WT - dox
WT + dox
D
α-catenin
α18
E
Nuclei/FOV
***
F
G
0 h
4 h
8 h
12 h
H
6 h
12 h

**Supplementary figure 6. Tubule formation assay of SHANK3-depleted cells and tissue dynamics of wild type HUVECs**

**A.** Tubule formation assay of shSHANK3 HUVECs +/- doxycycline (dox) seeded onto Matrigel for 16 h. Scale bar 200 µm. **B.** Quantification of tubules from A using Angiogenesis Analyzer for ImageJ (Carpentier et al., 2020). n = 3 biological replicates WT cells and 2 biological replicates shSHANK3 cells, 2-3 wells imaged per replicate. One-way ANOVA with Šídák test for multiple comparisons used for statistical analysis. **C.** Immunofluorescence imaging of WT HUVECs +/- dox seeded onto Matrigel for 16 h. Scale bar 200 µm. **D.** Immunofluorescence images of WT HUVEC monolayers +/- dox stained for total α-catenin and force-sensitive α-catenin conformation (α18). Scale bar 20 µm. **E.** Quantification of number of nuclei per field of view of cells in Supplementary Fig. 6D and Fig. 5D. n = 4 biological replicates, 9-11 fields of view quantified per replicate. One-way ANOVA with Šídák test for multiple comparisons used for statistical analysis. **F.** Quantification of nuclei per field of view in shSHANK3 HUVECs +/- dox imaged for traction force microscopy in Fig. 5F and G. n = 27-29 fields of view per condition across two independent experiments. Welch's t test used for statistical analysis. **G.** Spheroid wetting assay tracking WT HUVEC +/- dox migration away from spheroids over 12 h. Overlay indicates cell periphery at indicated time points. Scale bar 200 µm. **H.** Spheroid merge assay showing merging of WT HUVEC spheroids +/- dox over 12 h. Scale bar 100 µm. ns, non-significant; ***, p value < 0.005.

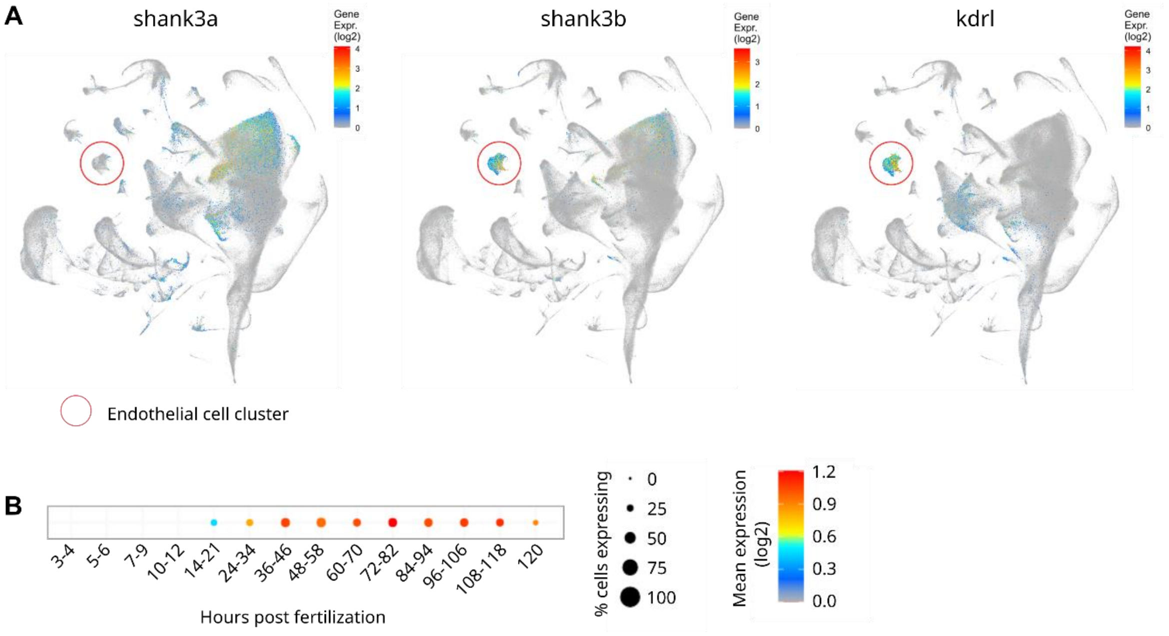


**Supplementary figure 7. *Shank3* expression in zebrafish**

**A.** Single-cell gene expression UMAP of *shank3* homologs, *shank3a* and *shank3b* and endothelial cell marker *kdrl* in zebrafish, showing expression of *shank3b*, but not *shank3a* in *kdrl* positive cells. **B.** Expression of *shank3b* in zebrafish embryo vasculature at varied timepoints post-fertilization. Data from Daniocell online database (Sur et al., 2023).

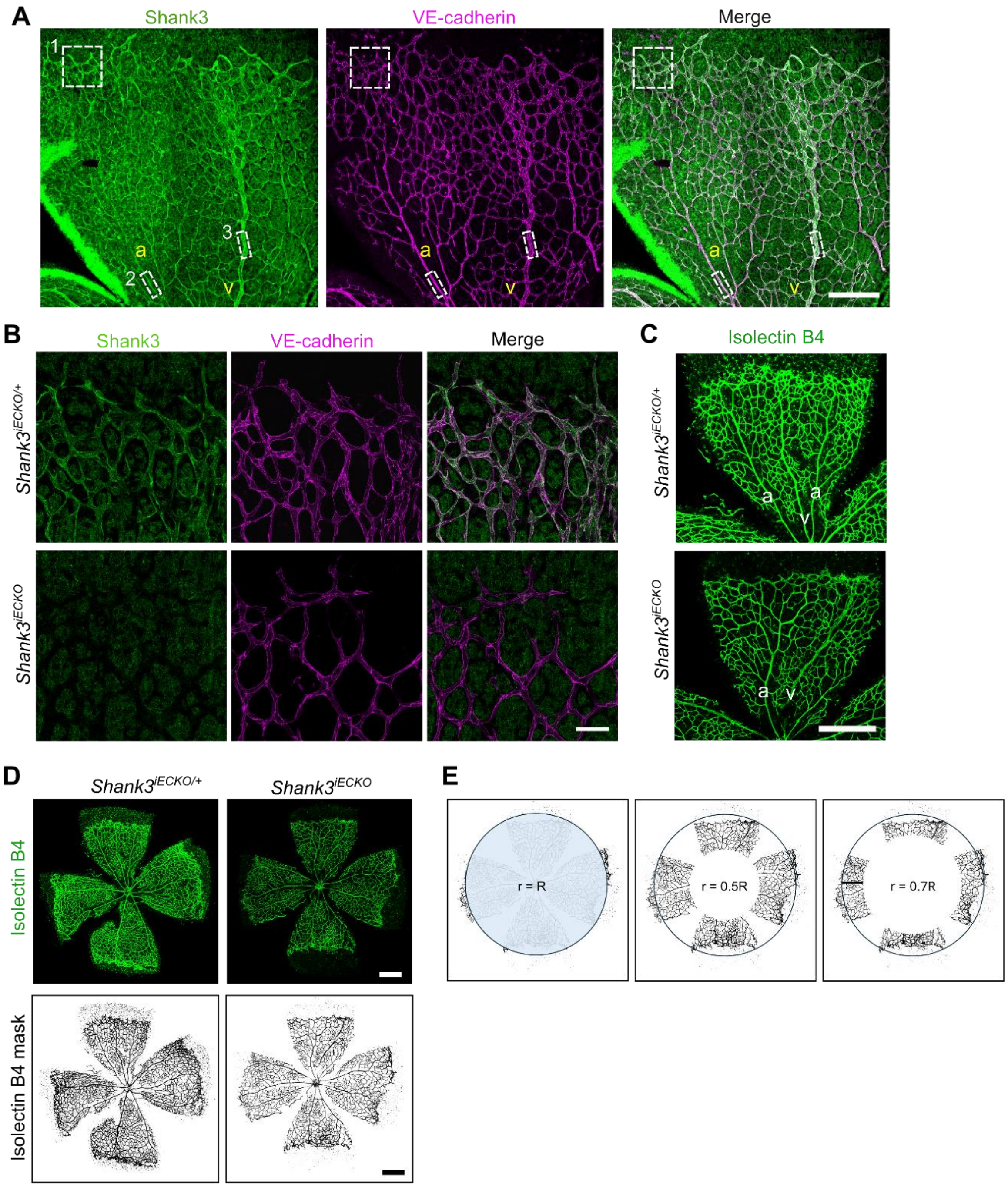


**Supplementary Figure 8. Analysis of vascular sprouts in *Shank3*$^{iECKO}$ retina at P6.**

**A.** Retinal blood vessels from P6 mice were stained for Shank3 and VE-cadherin. Inserts 1-3 indicate regions shown in Fig. 7A-C representing leading front (1), artery (2), and vein (3), respectively. Scale bar 200 µm. **B.** *Shank3*$^{flox/flox}$;tdTomato;Cdh5-CreERT2 mice (*Shank3*$^{iECKO}$) and *Shank3*$^{flox/+}$;tdTomato;Cdh5-CreERT2 littermates (*Shank3*$^{iECKO/+}$; controls) were administered 4-hydroxytamoxifen (4-OHT) at postnatal days 2–3 (P2–P3), followed by analysis of retinal vasculature at P6 (n = 4–5 mice per group). Representative images of Shank3 and VE-cadherin immunostaining in the retinal vasculature demonstrate loss of endothelial Shank3 expression in *Shank3*$^{iECKO}$ mice. Scale bar 50 µm. **C, D.** Representative images of isolectin B4 stained flat mounted retinas from *Shank3*$^{iECKO}$ and *Shank3*$^{iECKO/+}$ pups at P6. Mask from C is shown in Fig. 7F. Scale bar 500 µm. **C.** Representation of the proximal and distal zones of the retinal vasculature with respect to optic nerve head used for quantification in Fig. 7 (r = 0.5R and r = 0.7R). a, artery; v, vein.
